# The RNA binding proteins hnRNP H and F regulate splicing of a MYC dependent HRAS exon in Prostate Cancer Cells

**DOI:** 10.1101/2022.11.29.518269

**Authors:** Xinyuan Chen, Harry Taegyun Yang, Beatrice Zhang, John W. Phillips, Donghui Cheng, Frank Rigo, Owen N. Witte, Yi Xing, Douglas L. Black

**Affiliations:** Molecular Biology Interdepartmental Doctoral Program, University of California, Los Angeles, Los Angeles, California 90095, USA; Bioinformatics Interdepartmental Graduate Program, University of California, Los Angeles, California 90095, USA; Center for Computational and Genomic Medicine, The Children’s Hospital of Philadelphia, Philadelphia, Pennsylvania 19104, USA; Department of Microbiology, Immunology, and Molecular Genetics, University of California, Los Angeles, Los Angeles, California 90095, USA; Ionis Pharmaceuticals, Inc., 2855 Gazelle Ct., Carlsbad, California 92010, USA; Department of Molecular and Medical Pharmacology, University of California, Los Angeles, California 90095, USA; Jonsson Comprehensive Cancer Center, University of California, Los Angeles, California 90095, USA; Eli and Edythe Broad Center of Regenerative Medicine and Stem Cell Research, University of California, Los Angeles, California 90095, USA.; Molecular Biology Institute, University of California, Los Angeles, California 90095, USA; Department of Pathology and Laboratory Medicine, University of Pennsylvania, Philadelphia, Pennsylvania 19104, USA

**Keywords:** Alternative pre-mRNA splicing, post-transcriptional regulation, prostate cancer, MYC, HRAS, hnRNP H, hnRNP F, antisense oligonucleotides

## Abstract

The Myc proto-oncogene contributes to the pathogenesis of more than half of human cancers. Malignant transformation by Myc transcriptionally upregulates the core pre-mRNA splicing machinery and causes mis-regulation of alternative splicing. However, our understanding of how splicing changes are directed by Myc is limited. We performed a signaling pathway-guided splicing analysis to identify Myc dependent splicing events. These included an HRAS cassette exon repressed by Myc across multiple tumor types. To molecularly dissect the regulation of this HRAS exon, we used antisense oligonucleotide tiling to identify splicing enhancers and silencers in its flanking introns. RNA binding motif prediction indicated multiple binding sites for hnRNP H and hnRNP F within these cis-regulatory elements. Using siRNA knockdown and cDNA expression, we found that both hnRNP H and F activate the HRAS cassette exon.

Mutagenesis and targeted RNA immunoprecipitation implicate two downstream G-rich elements in this splicing activation. Analyses of ENCODE RNA-seq datasets confirmed hnRNP H regulation of HRAS splicing. Analyses of RNA-seq datasets across multiple cancers showed a negative correlation of hnRNP H gene expression with Myc hallmark enrichment, consistent with the effect of hnRNP H on HRAS splicing. Interestingly, hnRNP F expression showed a positive correlation with Myc hallmarks and thus was not consistent with the observed effects of hnRNP F. Loss of hnRNP H/F altered cell cycle progression and induced apoptosis in the PC3 prostate cancer cell line. Collectively, our results reveal new mechanisms for Myc-dependent regulation of splicing, and point to new possible therapeutic targets in prostate cancers.

**SIGNIFICANCE STATMENT:** Myc Transformation by the proto-oncogene c-Myc causes dysregulation of the pre-mRNA splicing reaction in cancer, but it is not known how mRNA isoform changes are directed by Myc. Here, we use bioinformatics to identify a splicing event in another proto-oncogene, HRAS, that is regulated by Myc across multiple tumor types. We identify new splicing regulators, hnRNP’s H and F, that control this HRAS exon by binding to enhancer elements within its downstream intron. Additional pan-cancer bioinformatic analyses show hnRNP H expression to be anti-correlated with Myc hallmarks, consistent with the reduced splicing of the HRAS exon in Myc driven cancer. These findings uncover new mechanisms by which Myc can alter splicing in cancer cells and provide new molecular targets for potential therapeutics.

## INTRODUCTION

Changes in pre-mRNA splicing have emerged as important contributors to the cancer phenotype. Core splicing components including the U snRNPs assemble onto nascent RNAs to form the catalytic spliceosome that will excise each intron (Wahl et al., 2009; Wilkinson et al., 2020). This assembly is regulated by RNA binding proteins that bind to the pre-mRNA at cis-regulatory elements to direct splice site choices and create alternatively spliced mRNA isoforms (Black, 2003; Fu and Ares, 2014; Lee and Rio, 2015). These regulators of splicing are very diverse and each alternative splicing event is regulated by multiple factors that can act either positively or negatively on the selection of a particular isoform. Aberrant splicing in cancer can result from mutations in core spliceosomal components that give rise to aberrant mRNAs, or from altered expression and modulation of regulatory RNA binding proteins that shift the production of particular mRNA isoforms (Dvinge et al., 2016; Escobar-Hoyos et al., 2019).

These changes in isoforms can affect many aspects of the tumor phenotype including cellular growth control and cell cycle progression, suppression of apoptosis, response to hormones and growth factors, loss of cellular differentiation, metastasis, angiogenesis and drug resistance (Urbanski et al., 2018). Splicing in tumor cells also appears to be more error prone, producing mRNAs that are not normally produced elsewhere and which provide appealing targets for immunotherapies (Pan et al., 2021; Smith et al., 2019; Wang and Aifantis, 2020).

Deregulation of the Myc proto-oncogene contributes to many cancers. Myc is a DNA binding protein that interacts across the genome resulting in a broad deregulation of transcription, including of genes encoding components of the core splicing machinery. These changes in the levels of spliceosomal components drive cancer associated changes in splicing, and Myc transformed cells have been shown to be unusually sensitive to splicing inhibition (Hsu et al., 2015; Koh et al., 2015). Myc transformation also enhances expression of multiple RNA binding regulators of splicing, leading to cancer associated changes in alternative splicing programs (Escobar-Hoyos et al., 2019). In prostate cancer, changes in the expression and function of multiple splicing regulators, including SAM68, hnRNP L, and others have been shown to contribute to the cellular phenotype (Caggiano et al., 2019; Fei et al., 2017; Ho et al., 2021).

One family of RNA binding proteins implicated in a variety of aspects of cancer are the heterogenous nuclear ribonucleoproteins H and F (hnRNP H/F). HnRNP H is encoded on three genes H1, H2, H3, and hnRNP F on one gene. HnRNPs H and F bind to G-run motifs GGG and GGGG, that act to enhance splicing of an alternative exon when present downstream and to repress splicing when present upstream or within the exon (Caputi and Zahler, 2001; Carlo et al., 1996; Chou et al., 1999; Garneau et al., 2005; Martinez-Contreras et al., 2006; Matunis et al., 1994; Min et al., 1995; Modafferi and Black, 1999, 1997; Schaub et al., 2007; Xiao et al., 2009). Activation of splicing by G-run elements is strongly affected by regulatory elements in the upstream exon and its 3’ splice site, indicating that hnRNP H/F activity requires additional cofactors (Modafferi and Black, 1999). The two proteins have similar effects on splicing, but slightly different binding specificities and can differ in their activities on particular target exons (Caputi and Zahler, 2001; Huelga et al., 2012). They can also form a heterodimer that may allow them to coordinately affect some targets (Min et al., 1995; Chou et al., 1999; Schaub et al., 2007). HnRNP H was found to be upregulated in glioma (LeFave et al., 2011), as well as colon cancer, and head and neck cancers (Rauch et al., 2010, 2011). Oncogenic splicing switches driven by hnRNP H include targets such as IG20/MADD in glioma (LeFave et al., 2011), TCF3 in lymphoma (Yamazaki et al., 2018), Mcl-1 and HER2 in breast cancer (Gautrey et al., 2015; Tyson-Capper and Gautrey, 2018), KHK in hepatocellular carcinoma (Li et al., 2016), and A-Raf in colon and head and neck cancers (Rauch et al., 2010). hnRNP H also regulates alternative splicing of the oncogenic fusion transcript EWS-FLI1 (Vo et al., 2022), and the RON protooncogene (Braun et al., 2018), and may alter translation in glioblastoma (Herviou et al., 2020). HnRNP F is less studied in the context of cancer cells but has been shown to be needed for the productive splicing of Sam68 in prostate cancer (Caggiano et al., 2019).

The availability of whole-transcriptome sequencing data across cancers has enabled the definition of splicing signatures in cancer tissues compared to normal cells. In an earlier study, we developed a pathway-guided transcriptomic analysis of prostate cancer using 876 RNA-seq datasets from cells ranging from normal prostatic tissue to aggressive prostate cancer (Phillips et al., 2020). This identified a program of 1039 cassette exons whose splicing correlated with Myc signaling during cancer progression. Myc-correlated exons were enriched in genes encoding splicing regulatory proteins and core spliceosomal components, as well as other cellular functions. The splicing of HRAS exon 5 was found to be particularly responsive to Myc activity, and the correlation between HRAS splicing and Myc activation is found in other tumor types (Phillips et al., 2020; Urbanski et al., 2022). HRAS belongs to the Ras oncogene family, regulates cell division, and is involved in multiple signal transduction pathways (Crespo and León, 2000; Moore et al., 2020). HRAS exon 5 affects overall expression from the gene, such that its inclusion leads to premature translation termination, and nonsense-mediated decay of the HRAS mRNA (Cohen et al., 1989). Transcripts that escape NMD encode a C-terminal truncated p19 Ras protein with distinct functions from the canonical p21 Ras protein (Guil et al., 2003a; Camats et al., 2009). High Myc levels lead to reduced exon 5 splicing and potentially higher levels of p21 HRAS protein.

Here we report that large scale bioinformatic analyses of splicing and Myc expression confirms the correlation of HRAS exon 5 repression with Myc signature score in prostate cancer and across many tumor types. To obtain mechanistic links between Myc oncogenic transformation and splice isoform choices we dissected the regulation of HRAS exon 5 splicing. We utilized antisense oligonucleotide tiling to identify intronic splicing enhancers and silencers adjacent to the exon. RNA binding motif enrichments indicated the presence of many hnRNP H/F binding sites within these cis-regulatory regions. We found that both hnRNP H and F activate HRAS exon 5 splicing and this activation required G4 and G3 elements in the downstream intron. Bioinformatic analyses of ENCODE RNA-seq datasets confirmed hnRNP H regulation of HRAS exon 5 and indicated that it is one of many exons regulated by both MYC and H/F. Additional pan-cancer bioinformatic analyses correlate the downregulation of hnRNP H expression and the upregulation of hnRNP F with Myc hallmark score. Loss of hnRNP H/F resulted in cell cycle arrest and induced apoptosis in prostate cancer cell lines. Taken together, our results reveal new mechanisms by which Myc alters splicing regulation and the phenotype of cancer cells.

## RESULTS

### Pathway Enrichment-Guided Activity Study of Alternative Splicing (PEGASAS) identifies HRAS exon 5 as repressed by Myc transformation across multiple tumor types

The protooncogene HRAS contains a conserved poison exon (exon 5) that alters both its expression and function. The exon-skipped isoform encodes the full length functional p21 HRAS protein, while the exon-included isoform contains a premature termination codon (PTC) that triggers the nonsense-mediated decay (NMD) of the HRAS transcripts. Transcripts that escape from NMD are translated into a C-terminal truncated p19 HRAS protein (Cohen et al., 1989; Guil et al., 2003a) (**Figure 1A**). HRAS p21 and p19 share most of the N-terminal G domain that mediates GTP hydrolysis (Pálfy et al., 2020; Simanshu et al., 2017). However, p19 lacks the last 16 amino acids of the allosteric lobe, and is reported not to bind GTP (Guil et al., 2003a) (**Figure 1B**). HRAS p21 also has a C-terminal hypervariable region (HVR) that is responsible for membrane binding and trafficking (Simanshu et al., 2017). HRAS p19 replaces this C-terminal domain with a C-terminal 20 amino acid sequence that is conserved across species, but whose function is not known. Several bioinformatic studies have connected HRAS exon 5 splicing with Myc transformation. HRAS exon 5 inclusion was found to be anti-correlated with Myc activity across prostate and breast cancers (Phillips et al., 2020). Greater skipping of this exon was also seen in Myc-active tumors in a pan-cancer analysis that implicated a network of SR proteins in its regulation (Urbanski et al., 2022).

**Figure 1.**
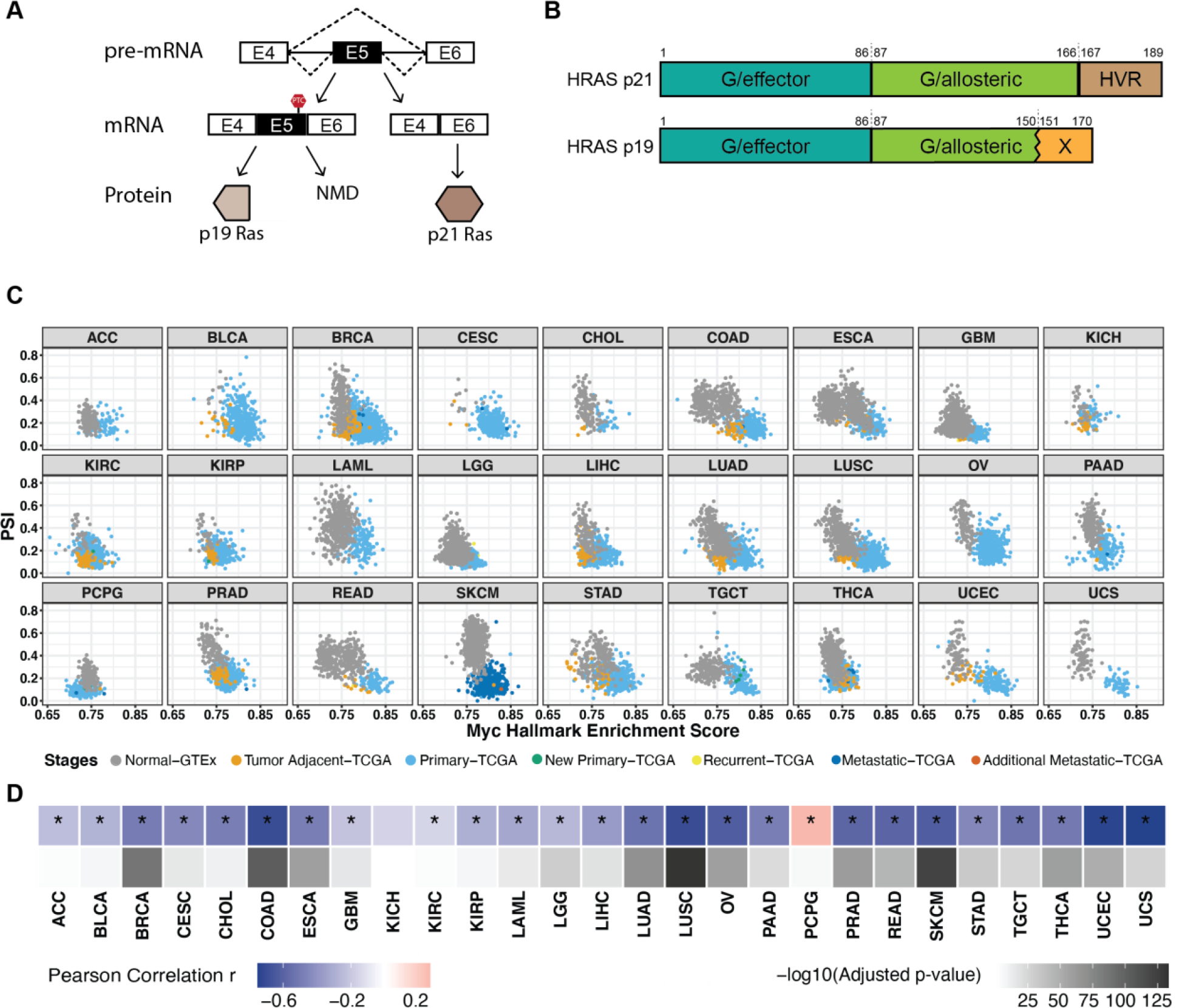
Pan-cancer analysis indicates that HRAS exon 5 is repressed by Myc activation cross multiple tumor types. **(A)** Diagram of HRAS pre-mRNA alternative splicing. NMD, nonsense-mediated decay. **(B)** Domain diagram of p21 and p19 HRAS isoforms (Pálfy et al., 2020). G/effector, G domain/effector lobe; G/allosteric, G domain/allosteric lobe; HVR, hypervariable region. **(C)** Scatterplot matrix showing the correlation of HRAS exon 5 PSI with Myc hallmark enrichment score across multiple tumor types. ACC, Adrenocortical carcinoma; BLCA, Bladder Urothelial Carcinoma; BRCA, Breast invasive carcinoma; CESC, Cervical squamous cell carcinoma and endocervical adenocarcinoma; CHOL, Cholangiocarcinoma; COAD, Colon adenocarcinoma; ESCA, Esophageal carcinoma; GBM, Glioblastoma multiforme; KICH, Kidney Chromophobe; KIRC, Kidney renal clear cell carcinoma; KIRP, Kidney renal papillary cell carcinoma; LAML, Acute Myeloid Leukemia; LGG, Brain Lower Grade Glioma; LIHC, Liver hepatocellular carcinoma; LUAD, Lung adenocarcinoma; LUSC, Lung squamous cell carcinoma; OV, Ovarian serous cystadenocarcinoma; PAAD, Pancreatic adenocarcinoma; PCPG, Pheochromocytoma and Paraganglioma; PRAD, Prostate adenocarcinoma; READ, Rectum adenocarcinoma; SKCM, Skin Cutaneous Melanoma; STAD, Stomach adenocarcinoma; TGCT, Testicular Germ Cell Tumors; THCA, Thyroid carcinoma; UCEC, Uterine Corpus Endometrial Carcinoma; UCS, Uterine Carcinosarcoma. **(D)** Heatmap summarizing the Pearson correlation coefficient for PSI vs Myc score in each tumor type, accompanied with corresponding adjusted p-value. *, tumor types with statistically significant adjusted p-value (< 0.05).

To broadly assess HRAS splicing changes in response to Myc signaling pathway activation in tumors, we used the computational framework PEGASAS (Phillips et al., 2020) to analyze RNA-seq data compiled from 9,490 samples from The Cancer Genome Atlas (TCGA) and 5,862 tissue matched samples from the Genotype-Tissue Expression project (GTEx) (Lonsdale et al., 2013; Weinstein et al., 2013). Briefly, gene expression and exon inclusion (Percent spliced in, PSI) values were computed for all genes and exons for each sample. Myc activity scores were calculated using the Myc Targets V2 hallmark gene set from the Molecular Signatures Database (MSigDB) (Liberzon et al., 2015). Myc activity scores were then correlated with all exon PSI values across all the datasets. Myc activity was seen to increase with disease progression from normal tissue (gray), to tumor adjacent benign tissue (orange), and to primary tumor (light blue) and more malignant disease stages. Inclusion of HRAS exon 5 is found to negatively correlate with Myc hallmark enrichment in the majority of 27 tumor types (**Figure 1C**). In addition to the correlation that we observed previously in prostate adenocarcinoma (PRAD), other epithelial cell cancers, such as colon adenocarcinoma (COAD) and lung squamous cell carcinoma (LUSC) showed particularly strong correlations (**Figure 1D**).

### ASO tiling reveals splicing enhancers and silencers controlling HRAS exon 5

As a first step in delineating the regulatory elements in the region of exon 5, we applied antisense oligonucleotides (ASOs) that base pair and potentially block RNA regulatory elements (Crooke et al., 2021). We designed and synthesized 22 ASOs that tiled across the highly conserved sequences of the HRAS exon 5 region, including 94 nucleotides of intron 4, the 82nt exon, and 200nt of intron 5 (**Figure 2A**). These ASOs were 18nt in length, non-overlapping, and had a uniform phosphorothioate backbone chemistry with methoxyethyl modifications at the 2’ ribose positions (2’MOE-PS). Each ASO, along with a non-targeting control (NTC) ASO, was transfected into HEK293 cells. 24 hours after transfection, RNA was isolated and HRAS splicing was measured by semi-quantitative RT-PCR. Comparing the exon 5 percent-spliced in (PSI) value in the presence of each ASO to the NTC, we identified ASO’s that decrease exon 5 splicing and others that increase it (**Figure 2B**). As expected, ASOs targeting the 3’ and 5’ splice sites strongly inhibited splicing. ASOs targeting the body of the exon also induced exon skipping, indicating the presence of exonic splicing enhancers within the HRAS exon. ASOs targeting downstream intron 5 showed diverse effects with some increasing and others decreasing exon 5 splicing, suggesting the presence of multiple intronic splicing silencers and enhancers. ASOs I5-1, I5-4, and I5-7 increased exon inclusion and may block intronic splicing silencer elements (ISSs). In contrast, ASOs I5-3, I5-5, I5-9, and I5-11 all induced exon skipping compared to controls and suggest the presence of splicing enhancers in this region.

**Figure 2.**
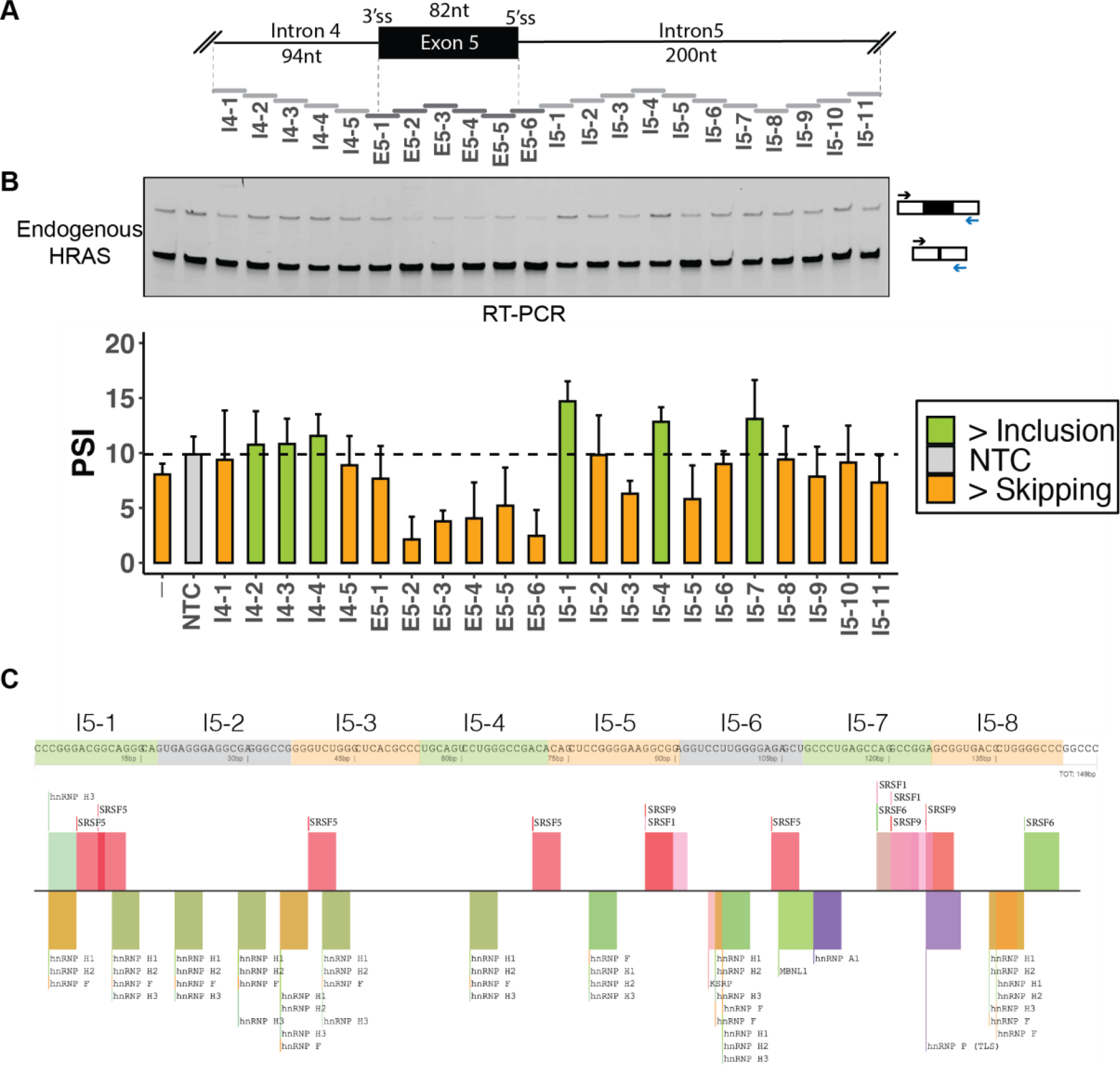
Identification of splicing cis-regulatory elements controlling HRAS exon 5. **(A)** Schematic of ASO tiling across HRAS exon 5 and its partial flanking introns, including 376- nucleotide region tiled by non-overlapping 18-mer ASOs. Each horizontal bar represents an ASO. **(B)** Semi-quantitative RT-PCR analyses showing the effects of ASOs on endogenous HRAS splicing. The arrows indicate RT-PCR primers for assay of exon 5 in the endogenous HRAS transcripts. The bar graph presents the quantification of the RT-PCR calculated as percent-spliced in (PSI) (gray: control; orange: more skipping; green: more inclusion). Each bar represents the mean value +/- SD of three biological replicates. NTC, non-targeting control. -, no-ASO mock control. **(C)** SpliceAid2 snapshot of predicted splicing factor binding sites on HRAS intron 5. The sequences targeted by each ASO are highlighted in orange (more skipping), green (more inclusion), or gray (no change).

We next constructed an HRAS minigene reporter by cloning the genomic region spanning exon 5, including the flanking introns and portions of exons 4 and 6, into the pcDNA3.1(+) expression vector (**Figure S1A**). We introduced an in-frame ATG start codon downstream from the CMV promoter, and a TGA stop codon upstream of the BgH polyadenylation site to reduce nonsense mediated decay of the product mRNA. In HEK293 cells, transcripts from the minigene show 18.9% exon 5 inclusion compared to 8.0% inclusion in the endogenous HRAS transcripts (**Figure S1B-C**). In co-transfection experiments we found that ASOs had pronounced effects on splicing the HRAS minigene. With some exceptions, the ASO induced changes in minigene splicing were consistent with those on endogenous HRAS transcripts. ASO’s targeting the exon were strongly inhibitory. Enhanced splicing, indicating the position of an intronic splicing silencer, was seen with I5-1 and I5-7. The adjacent I5-6 ASO also enhanced splicing of the minigene transcripts, but had more limited effect on the endogenous transcripts. Splicing inhibition due to the likely blocking of splicing enhancers was seen with ASO’s I5-5, and I5-8 to I5-11, often with stronger effects on the minigene RNA than seen on the endogenous RNA (**Figure 2B, S1C**). ASO I4-1 in intron 4 generated a new band in the RT-PCR likely due to a cryptic splice site in the minigene construct. The weaker effects of the ASO’s on the endogenous RNA may result from a number of factors. Different rates of transcription for the native and transfected genes could affect the ability of ASO’s to act on the RNA. Also mature endogenous RNA is already present at the time of ASO transfection and may not turn over completely during the assay of the ASO’s. In contrast, the minigene is co-transfected with the ASO’s so all RNA is processed in their presence. A previous study identified a splicing silencer element called rasISS1 in HRAS intron 5 (Guil et al., 2003b) (**Figure 4A**). The inhibitory sequence of rasISS1 maps to the region covered by ASOs I5-1 and I5-2. Our data also indicate a silencer in the I5-1 region. The limited effect we observe from I5-2 may be due to the secondary structure proposed for this region interfering with targeting by antisense oligos (Guil et al., 2003b). We have focused on regulatory elements where ASO’s inhibited splicing of the endogenous HRAS exon indicating intronic enhancer elements (I5-3, I5-5, and I5-8).

To identify trans-acting splicing factors that potentially bind the HRAS splicing regulatory elements, we examined the sequences surrounding HRAS exon 5 with motif finding tools SpliceAid2 and RBPmap (Paz et al., 2014; Piva et al., 2012). This identified many binding motifs for hnRNP H and its paralog hnRNP F downstream of exon 5, as well as motifs for hnRNP A1, SRSF5, and other proteins (**Figure 2C, S2A**). Previous work identified hnRNP A1 as repressor of exon 5 that binds to the rasISS1 element (Guil et al., 2003b). The SR proteins SRSF2 and SRSF5 were also identified as factors that correlate with exon 5 activation (Guil et al., 2003b).

### hnRNPs H and F activate HRAS exon splicing

Heterogenous nuclear ribonucleoproteins H and F (H1, H2, H3 and F) are paralogous splicing factors that activate splicing when bound downstream of alternative exons. H and F both bind to G-run motifs GGG and GGGG, although they exhibit slightly different binding specificities (Caputi and Zahler, 2001; Chou et al., 1999; Garneau et al., 2005; Martinez-Contreras et al., 2006; Matunis et al., 1994; Min et al., 1995; Schaub et al., 2007). HnRNP H3 is a pseudogene that lacks the N-terminal RNA Recognition motif (Mahé et al., 1997; Honoré, 2000). HnRNP’s H1 and H2 are highly similar in peptide sequence (Honoré et al., 1995), but differ in their relative expression, with H1 more highly expressed in most cell types including HEK293 cells and a Myc dependent prostate cancer model (Lonsdale et al., 2013; Phillips et al., 2020). (Honoré et al., 1995; Mahé et al., 1997). We have focused on hnRNP H1 and F.

To assess the effects of hnRNPs F and H on HRAS splicing, we performed siRNA-mediated knockdown of these factors individually and together in HEK293 cells. Immunoblot confirmed that the siRNAs depleted hnRNP F by 93% and H1 by 70% when they were targeted individually, and by 89% and 68% when hnRNPs F and H were targeted together (**Figure 3A**). Depletion of F or H individually led to a modest increase in the other factor, an apparent cross-regulation that is commonly observed for paralogous pairs of RNA binding proteins. 72 hours after introduction of the siRNAs, we assayed the splicing of exon 5 in the endogenous HRAS transcripts by RT-PCR. Depletion of either hnRNP F or hnRNP H decreased exon 5 splicing, with a stronger effect seen with the loss of hnRNP H, despite its less complete depletion.

**Figure 3.**
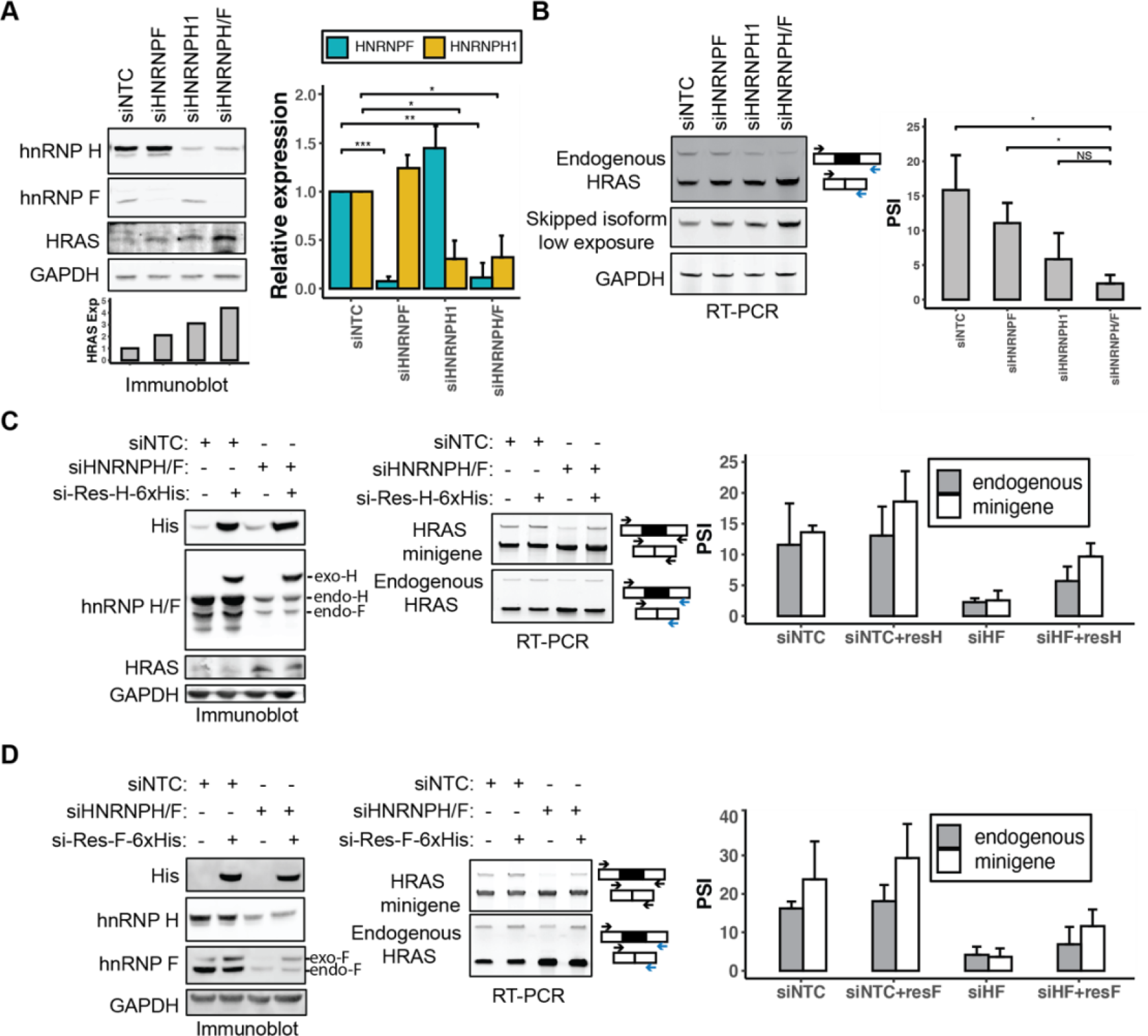
hnRNP H and F activate HRAS exon 5 splicing. **(A)** Immunoblot showing the expression of hnRNP H, hnRNP F, and HRAS proteins in HEK293 cells, after transfection with control, HNRNPF, HNRNPH1, or HNRNPH/F siRNAs. GAPDH was used as a loading control. The bar graph (right) shows the quantification of hnRNP H and F proteins in response to each siRNA perturbation. The grayscale bar graph (bottom) shows the quantification of HRAS p21 protein expression in response to each siRNA perturbation. NS: p > = 0.05; *: p < 0.05;**: p < 0.01; ***: p < 0.001 (Student’s t-test). **(B)** RT-PCR analysis endogenous HRAS splicing after siRNA knockdown of hnRNP’s H and F. The bar graph (right) shows the quantification of the RT-PCR results. **(C-D)** Rescue of hnRNP H/F expression after knockdown in HEK293 cells. Immunoblot of hnRNP H, hnRNP F, His tagged rescue protein, and HRAS in HEK293 cells. Cells were transfected with control or HNRNPH/F siRNAs followed by transfection with siRNA- resistant C-terminal 6xHis tagged hnRNP’s H (C) or F (D). RT-PCR of minigene and endogenous HRAS splicing in each experimental condition is quantified in the bar graphs (right). Each bar represents the mean +/- SD of three biological replicates.

Depletion of both H and F resulted in even greater exon skipping. Thus, both H and F act to enhance HRAS splicing (**Figure 3B**). H/F knockdown resulted in an upregulation of the HRAS p21 protein isoform as seen by immunoblot, consistent with the observed splicing changes (**Figure 3A**). To rule out off-target effects of the siRNAs, we re-expressed 6xHis tagged siRNA-resistant HNRNPH1 or HNRNPF cDNAs, after siRNA depletion of the endogenous transcripts (**Figure 3C-D**). Immunoblots confirmed the expression of recombinant hnRNP H or F at levels comparable to the endogenous proteins. Re-expression of either hnRNP H or F stimulated splicing of HRAS exon 5 in both minigene and endogenous RNAs, thus validating their roles as splicing activators and ruling out possible off target effects of the siRNAs.

### G_1_ and G_4_ elements in the downstream intronic splicing enhancer mediate the hnRNP H/F dependent enhancement of HRAS splicing

HnRNPs H and F bind to motifs containing runs of three or four G nucleotides which act as splicing enhancers when found in a downstream intron (Chou et al., 1999; Schaub et al., 2007). There are ten G-runs downstream of exon 5, of which four were identified by the ASO tiling as enhancer elements (**Figure 4A**). We constructed a series of minigene reporters carrying single mutations at each of these four G-run elements, a double mutation of the neighboring G1 and G2 runs, and a mutation of the four G-runs (G1,G2,G3,G4). After transfection into HEK293 cells, the splicing of each of these constructs was compared to the wildtype clone by RT-PCR (**Figure 4B**). We observed small decreases in exon 5 splicing when either G1 or G2 was mutated, and minimal splicing changes resulting from G3 or G4 mutations. In the constructs carrying single G-run mutations the expression of recombinant F or H enhanced exon inclusion. The double mutation of both G1 and G2 resulted in a nearly complete loss of exon 5 splicing and a similar effect was seen when all four G runs were mutated together. Notably, for these mtG1G2 and mtG1G2G3G4 constructs, the overexpression of H or F could no longer rescue exon 5 splicing. We also tested mutations in other G-runs downstream of exon 5 but where the blocking ASO either indicated the presence of a splicing silencer or had minimal effect (G-runs located in the I5-1, I5-4 and I5-6 targeted regions, Figure 4A). The splicing changes induced by these mutations were mostly consistent with the ASO data, except one G-run in the I5-1 region whose mutation indicated that it also acted as an enhancer but the ASO was apparently blocking a silencer (Supplementary table 5). This region is thus complex and likely contains multiple regulatory elements, one of which might be an additional hnRNP H/F dependent enhancer. Altogether the results indicate that multiple G-runs within HRAS intron 5 act as hnRNP H/F dependent splicing enhancers, with individual elements acting redundantly and with G1 and G2 having the strongest effects.

**Figure 4.**
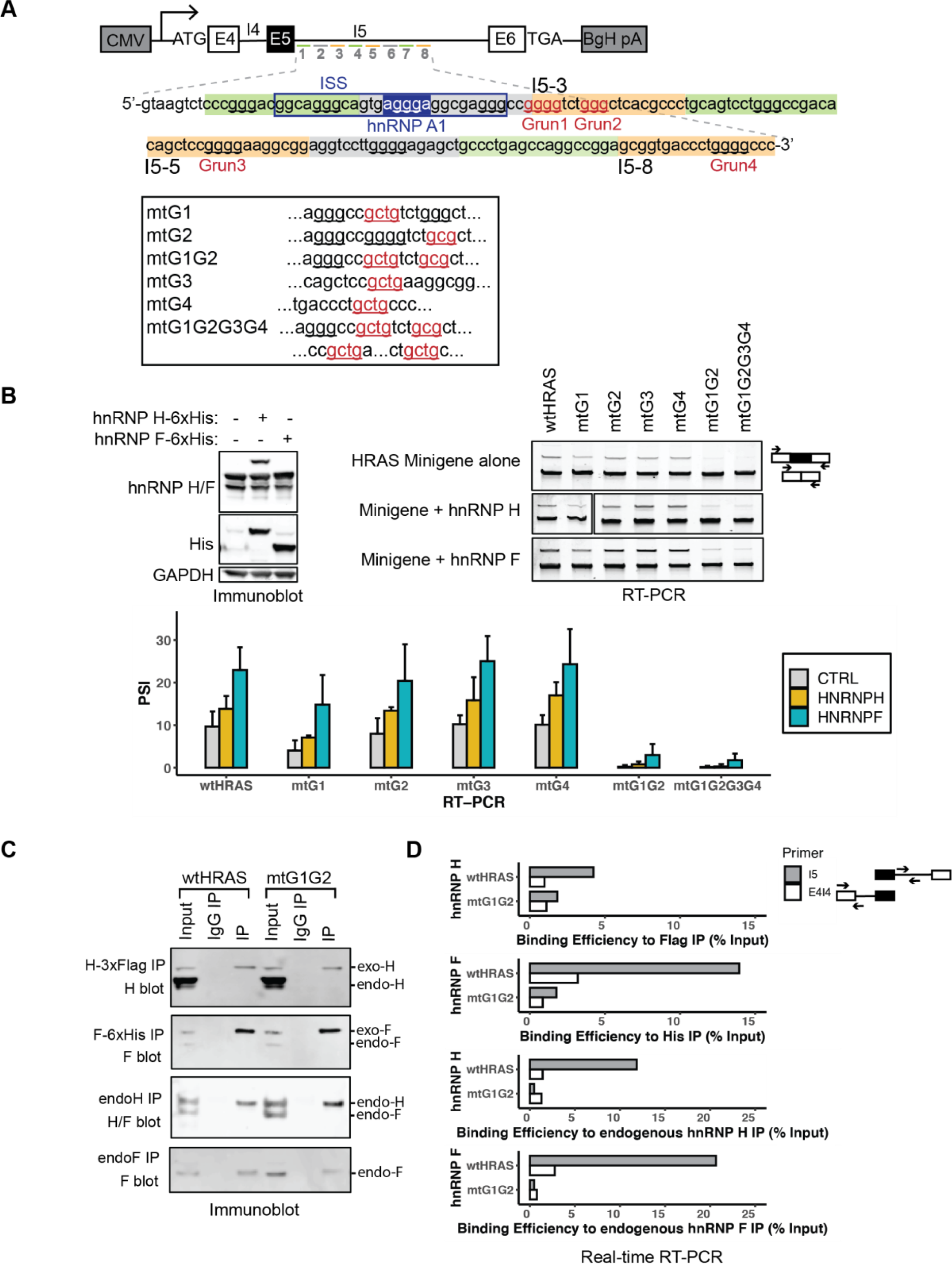
hnRNP H and hnRNP F modulate HRAS exon 5 splicing through G-run elements within the downstream ISE. **(A)** Diagram of HRAS minigene reporters carrying mutations at putative hnRNP H/F binding motifs. Intron 5 nucleotide sequences targeted by ASO’s I5-1 to I5-8 are shown in orange as potential enhancers and green as potential suppressers. G-runs within this intron 5 region is underlined, and those that are within the enhancer regions and mutated are labeled in red and numbered. Mutations in these elements are indicated below, with the mutated sequences underlined and labeled in red. The known silencer element rasISS1 is boxed in navy and its putative hnRNP A1 binding motif is highlighted (Guil et al., 2003b). **(B)**Immunoblot showing the expression of endogenous hnRNP’s H and F and His-tagged ectopic proteins. GAPDH was used as the loading control. RT-PCR analyses showing the splicing changes of wild type and mutant minigene reporters after co-transfection with control (top), hnRNP H (middle), and hnRNP F (bottom) expression plasmids (wt, wild type; mt, mutant). The bar graph presents the quantification of RT-PCR results, with the mean +/- SD of three biological replicates. **(C)**RNA immunoprecipitation assayed by real-time RT-PCR of wildtype and G1G2 mutant HRAS intron 5 RNA bound by hnRNP H and hnRNP F proteins. HEK293 cells transfected with wtHRAS or mtG1G2 minigenes were immunoprecipitated using anti-epitope tag (Flag or His) antibodies, anti-endogenous protein (hnRNP H or hnRNP F) antibodies, or non-immune IgG. The immunoblot shows the recovery of hnRNP H, hnRNP F or both in the IP’s. **(D)** The bar graphs (right) of real-time RT-PCR quantification of recovered RNA using two sets of primers: the I5 product targets an intronic region neighboring the mutation and the E4I4 product targets the sequence immediately upstream.

To assess the interactions of hnRNPs H and F with HRAS intron 5, we assayed for the presence of HRAS RNAs in hnRNP H/F immunoprecipitates. HEK293 cells were transfected with the wildtype HRAS minigene or that carrying the G1G2 mutation. These were examined in cells expressing 3xFlag tagged hnRNP H or 6xHis tagged hnRNP F, or in untransfected cells to assay endogenous H and F protein. Ectopically expressed H and F were immunoprecipitated with anti-Flag and anti-His antibodies respectively, and endogenous hnRNPs H and F with antibodies reactive with each protein. The specificity of the pull downs and yield of the immunoprecipitations were monitored by immunoblot (**Figure 4C**). RT-PCR analysis of the immunoprecipitated RNA identified HRAS pre-mRNA associated with the H and F immunoprecipitates but not the control IgG. Primer pairs were designed to amplify either the exon4-intron4 junction or a segment of intron 5 neighboring the G-runs. These yielded the expected products from the minigene reporter whose unspliced products were much more abundant than the nascent endogenous HRAS RNA. A minus reverse transcriptase control confirmed the absence of minigene DNA (**Figure S3A**). To quantify the amounts of mutant and wildtype RNAs in each pulldown, we performed qRT-PCR, normalizing the amount of RNA in the precipitates to the input. We found that the binding of HRAS RNA to hnRNPs H and F was substantially reduced by G1G2 mutation. This was seen in immunoprecipitates of both the exogenous and endogenous proteins (**Figure 4D**). The Intron 5 fragment was more abundant in the immunoprecipitates than the exon 4 - intron 4 fragment and showed the biggest reduction in binding with the G1G2 mutation. However, reduced binding was also observed with the exon-intron fragment. Taken together, the data indicate hnRNPs H and F interact with the G-runs in HRAS intron 5 to activate exon 5 splicing.

### HNRNPH gene expression is repressed by Myc activation across multiple tumor types

To further assess the association of hnRNPs H and F with Myc, we performed a correlation analysis of Myc activity score and normalized splicing factor expression across tumor types. We used DEseq2 to normalize the read counts of 220 genes encoding splicing factors (Han et al., 2013) in the TCGA and GTEx samples analyzed above. We then correlated splicing factor expression with Myc activity scores computed from PEGASAS. HNRNPH1 exhibited a clear negative correlation with Myc activation in the majority of 27 tumor types, including prostate cancer (PRAD, **Figure 5A-B**). In contrast, HNRNPF exhibited a positive correlation with Myc activation in almost all tumor types (**Figure S4A-B**). The negative correlation between HNRNPH1 and Myc was found in multiple epithelial cancers such as breast invasive carcinoma (BRCA), colon adenocarcinoma (COAD), lung squamous cell carcinoma (LUSC) and Ovarian serous cystadenocarcinoma (OV). Interestingly, the correlations were reversed in the acute myeloid leukemia (LAML) samples. In these tumors hnRNP H1 was positively correlated with Myc and hnRNP F negatively correlated.

**Figure 5.**
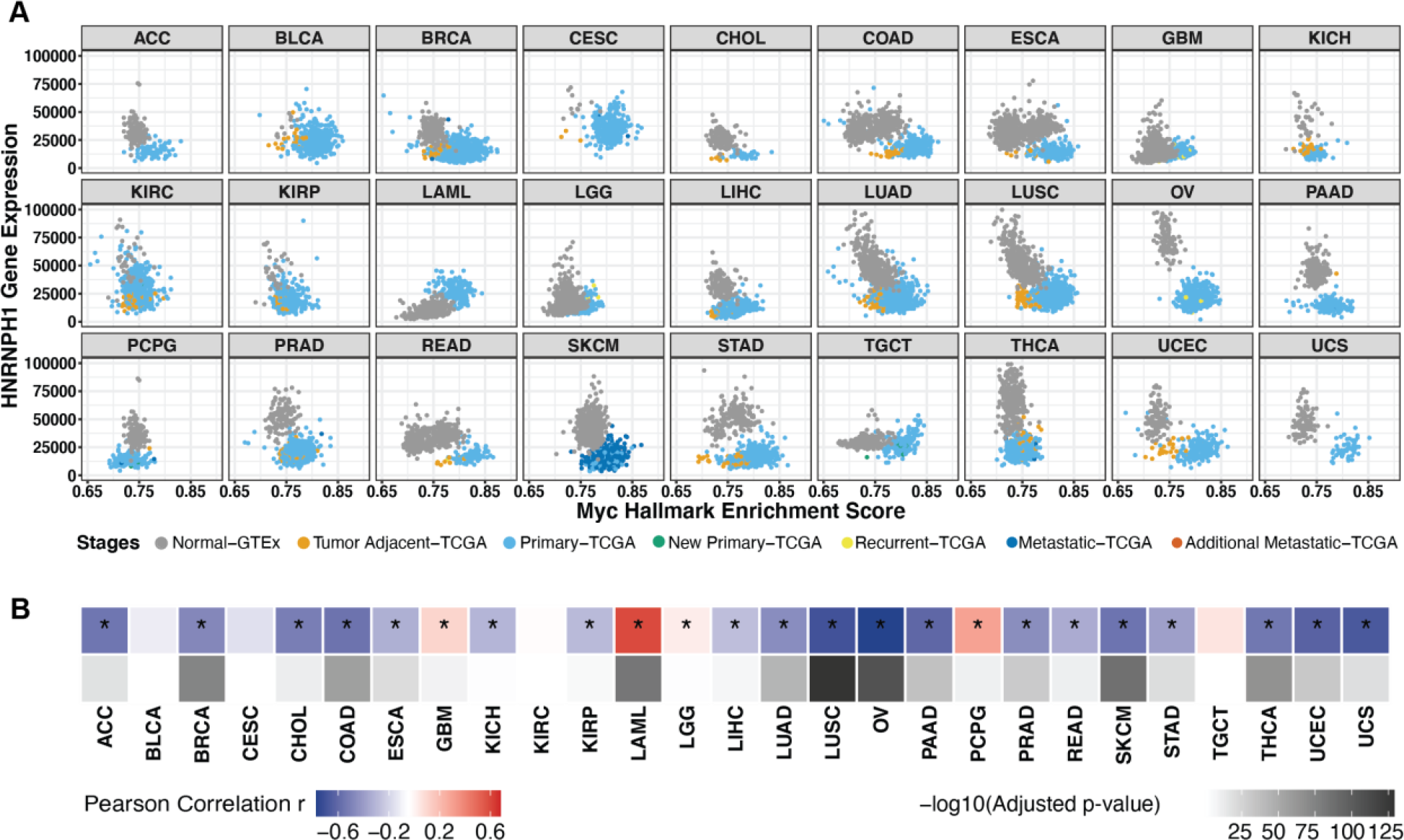
HNRNPH1 gene expression is downregulated with Myc activation across tumor types. **(A)** Scatterplot matrix showing the correlation of normalized HNRNPH1 versus Myc hallmark enrichment score across the disease spectrum in multiple tumor types. **(B)** Heatmap summarizing the Pearson correlation coefficient of Myc vs hnRNP H expression for each tumor type, accompanied with its adjusted p-value. *, tumor types with statistically significant adjusted p-value (< 0.05).

### The HRAS exon is one of many exons controlled by both MYC and hnRNP H/F activity

To identify additional exons regulated by hnRNPs H and F, we analyzed RNA-seq datasets from the ENCODE project (Dunham et al., 2012). Using rMATS-turbo, we compared RNA from HepG2 cells after HNRNPH or HNRNPF knockout to RNA from the non-targeted control cells (**Figure 6A, S5A**) (Shen et al., 2014). This identified 2190 and 1516 changes in skipped exons (SE) after H and F knockout respectively. HnRNP H can act either as a splicing repressor or activator depending on its binding location (Huelga et al., 2012). Consistent with this, we observed that approximately 50% of the exons showed reduced splicing upon hnRNP H knockout, indicating the protein acted to enhance their splicing. These included HRAS exon 5 which exhibited a PSI of 0.11 in the HNRNPH1 knockout samples, compared with 0.27 in the non-targeted control (**Figure 6B**). Similar overall results were obtained with the hnRNP F knockout, except that HRAS exon 5 showed no significant change upon hnRNP F knockout compared with the non-targeted control. This limited effect might result from the lower expression of hnRNP F yielding a smaller effect on the PSI after knockout (**Figure S5A-B**). To further validate the ENCODE findings, we knocked down hnRNPs H and F by siRNA in HepG2 cells and performed RT-PCR on the endogenous HRAS transcripts. As seen in the ENCODE RNA-seq data and in HEK293 cells (Figure 3), the loss of either H or F alone reduced HRAS exon inclusion in HepG2 cells, and the double knockdown had a stronger effect (**Figure S5C**). Interestingly, in these cells the double depletion of H and F also slightly reduced MYC expression as seen by immunoblot (see below). The data for HepG2 and HEK293 both indicate that hnRNP H1 is an activator of HRAS exon 5 splicing (Figure 3B).

**Figure 6.**
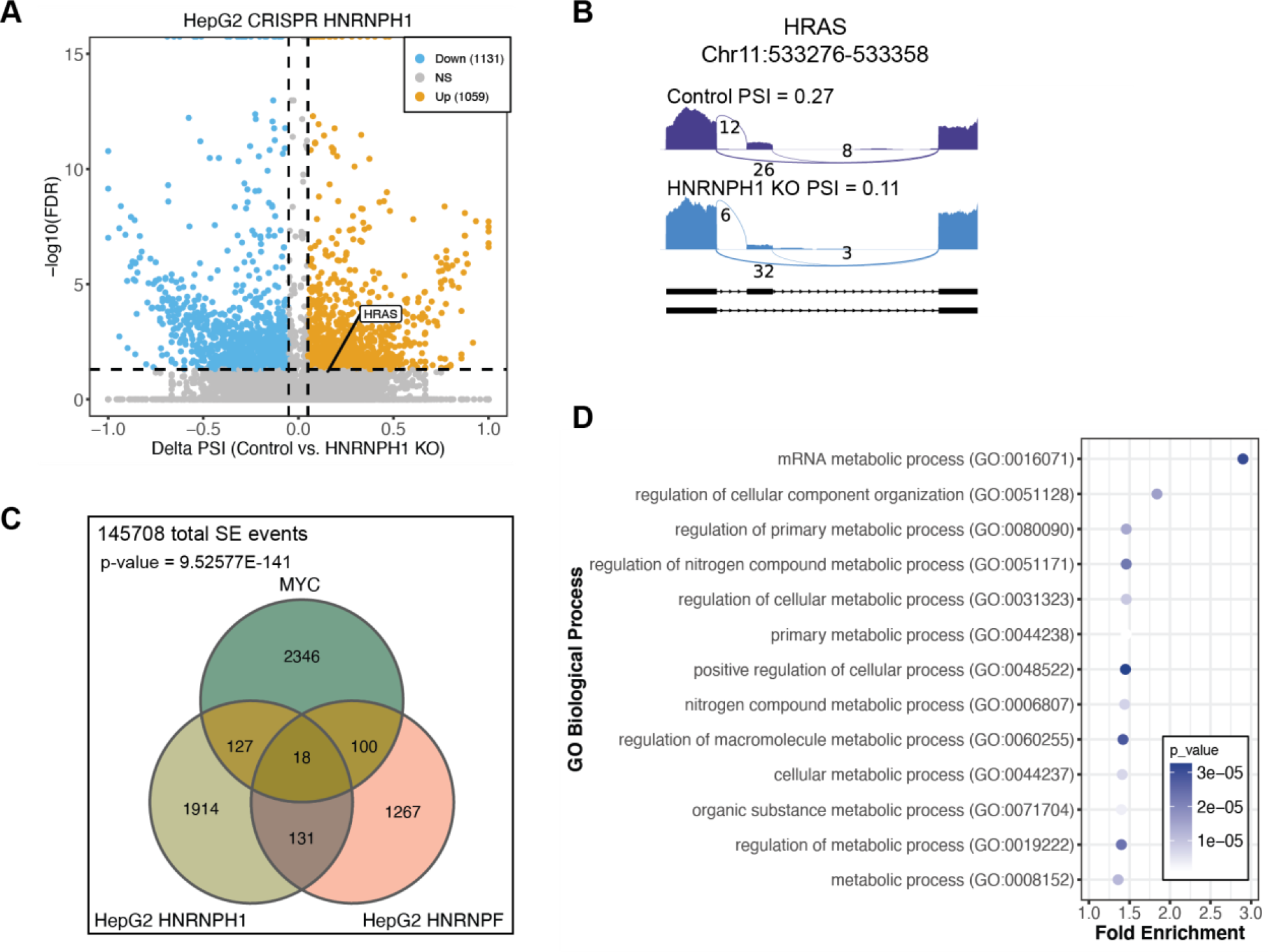
HRAS exon 5 is one of many exons controlled by both MYC and hnRNP’s H and F. **(A)** Scatterplot showing the SE (Skipped Exon) events detected by RNA-seq from the ENCODE project in HepG2 cells after HNRNPH1 CRISPR knockout. Significant SE events were filtered by junction reads per event > 10, |deltaPSI| > 0.05, and FDR < 0.05. **(B)** Sashimi plot showing the PSI of HRAS exon 5 in control cells and after HNRNPH1 KO. **(C)** Venn diagram showing the overlapping significant SE events across three datasets: MYC on vs. MYC off, HNRNPH1 KO vs Control, and HNRNPF KO vs Control. The numbers reflect overlapping events showing changes in each comparison without considering the direction of the changes. **(D)** Gene ontology analyses using PANTHER for genes represented by SE events between MYC and HNRNPH, between MYC and HNRNPF, or between both comparisons.

Analyses of the other ENCODE line, K562, also showed that HRAS exon 5 was positively regulated by hnRNP H1 (data not shown). However, in these cells hnRNP F gave different results exhibiting an increase in exon 5 splicing after hnRNP F knockout (data not shown). Presumably K562 expresses a different complement of splicing regulators that allow hnRNP F to act differently on some of its target RNAs. Notably, in our pan-cancer gene expression analysis (Figure 5, S4), we found both hnRNP H and F are under different transcriptional control by MYC in Acute Myeloid Leukemia (LAML), compared to most other tumor types. Thus, the lymphoblastic K562 cells derived from a chronic myelogenous leukemia may behave more similarly to LAML than to HepG2.

To examine whether other exons that are regulated by hnRNPs H and F are also regulated by Myc, similarly to HRAS, we analyzed RNA-seq data from a prostate cancer model carrying a doxycycline-inducible MYC gene (Phillips et al., 2020). This identified 2591 differentially spliced SE events between the MYC-on and MYC-off conditions. These MYC-dependent SE events overlapped with several hundred of the hnRNP H1 or F -dependent SE events from HepG2 (**Figure 6C**). A hypergeometric test comparing these exon sets yielded a p-value of 9.53xE^-141^ indicating a statistically significant fraction of the splicing events regulated by MYC are also controlled by hnRNP H1 and/or hnRNP F. Gene ontology analyses (PANTHER) (Thomas et al., 2022) indicate that the genes containing splicing events in the three way overlap are enriched for genes involved in metabolic processes, with mRNA metabolic processes as the top term (**Figure 6D**). Thus, there are multiple exons controlled by both MYC and hnRNPs H and F, and HRAS exon 5 is just one example.

### hnRNP H/F are required for cell proliferation in prostate cancer cell lines

To evaluate effects of splicing factors hnRNP H and F on the growth of Myc transformed cells, we knocked down their expression in two prostate cancer cell lines. PC3 is an advanced adenocarcinoma cell line with high metastatic potential, while DU145 cells derive from a prostate carcinoma with moderate metastatic potential (Namekawa et al., 2019). PC3 and DU145 cells transfected with siRNAs targeting hnRNP’s F, H, or both showed >75% depletion of each factor. The combined depletion of hnRNP H and F induced HRAS exon 5 skipping as seen previously (**Figure 7A-B**). The knockdown of H or F or both also resulted in a small reduction in MYC in the PC3 cells. The double knockdown also showed an increase in cPARP protein indicating the induction of cell apoptosis (**Figure 7A**), and it was apparent that the cultures had stopped proliferating after the hnRNP H/F depletion. To evaluate how loss of hnRNP H and F affected cell growth, we performed flow cytometry of propidium iodide stained PC3 cells. Depletion of hnRNP F or hnRNP H, and particularly the double knockdown, reduced the number of cells in the G1 phase and increased cells in G2 compared with control cells (**Figure 7C-D**), indicative of a mitotic block. HnRNP’s H and F are thus needed for proper mitotic progression in the prostate cancer cell lines.

**Figure 7.**
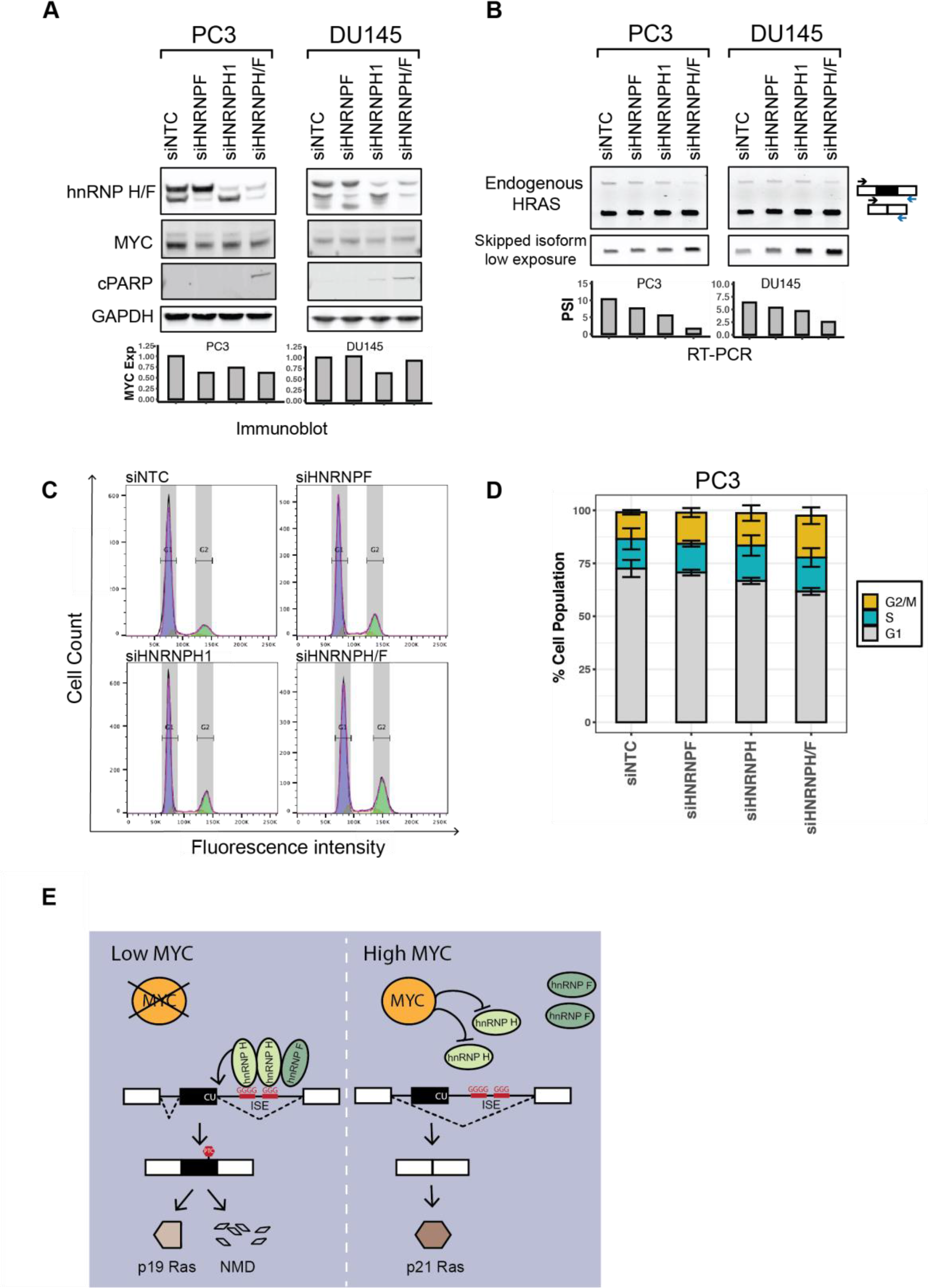
hnRNP’s H and F are required for cell proliferation in prostate cancer cell lines. **(A)** The effects of hnRNP H and hnRNP F knockdown on the expression of Myc and the apoptosis marker cPARP in the prostate cancer cell lines, PC3 and DU145. Immunoblot showing the expression levels of hnRNP’s H and F, MYC and cPARP in cells transfected with control, HNRNPF, HNRNPH1, or HNRNPH/F siRNAs. GAPDH was used as a loading control. The bar graphs present the quantification of MYC expression in response to each siRNA perturbation. **(B)** RT-PCR analysis showing the splicing changes of endogenous HRAS transcripts in response to each siRNA perturbation. The bar graphs show the quantification of HRAS exon 5 PSI. **(C)** Cell cycle analysis by FACS of PC3 cells transfected with control, HNRNPF, HNRNPH1, or HNRNPH/F siRNAs, and stained with propidium iodide. **(D)** Stacked bar plot showing the quantification of cells in the G1, S, and G2/M phases. Bars present the mean +/- SD of three biological replicates**. (E)** Model of hnRNP H regulation of HRAS exon alternative splicing under low MYC versus high MYC conditions.

## DISCUSSION

We found that Myc transformation influences the splicing of HRAS protooncogene transcripts to decrease exon 5 inclusion and allow greater production of the full-length isoform. We identified clusters of positive and negative splicing regulatory elements in the sequence encompassing exon 5, including splicing enhancers in the intron downstream. We show that these G-run enhancers bind to the splicing regulators hnRNP H and hnRNP F and are required for activating splicing of exon 5. HnRNP H1 expression was found to be anticorrelated with Myc score across many tumor types including lung, breast, and prostate, consistent with the repression of exon 5 by Myc. HnRNP F showed the opposite correlation. We found that HRAS exon 5 is one of a group of Myc regulated exons that are also regulated by hnRNP’s H and F, which are essential for the growth of prostate cancer cells. These results are summarized in Figure 7E, where the activation of MYC downregulates hnRNP H, leading to decreased HRAS exon 5 splicing and increased expression of the full-length p21 HRAS oncoprotein.

Human HRAS exon 5 was originally named IDX and shown to carry an in-frame premature termination codon (PTC) to create a truncated protein (Cohen et al., 1989). Human mutations that reduce IDX inclusion were shown to increase the activity of full length p21 HRAS (Cohen et al., 1989; Cohen and Levinson, 1988). HRAS exon 5-containing transcripts were later shown to undergo nonsense mediated mRNA decay (NMD) in the cytosol, with the undegraded portion found largely in the nucleus (Barbier et al., 2007). We found that cycloheximide treatment does increase this isoform consistent with its loss to NMD (data not shown). Thus, one role of the exon seems to be the modulation of p21 HRAS levels and activity.

HRAS transcripts that escape from NMD are potentially translated into the C-terminal truncated p19 HRAS protein, reported to have a distinct function from the full length p21 HRAS protein (Guil et al., 2003a; Camats et al., 2009). Exon 5 and its flanking introns are conserved in mammals, although the mouse gene does not contain the stop codon and the exon 5-included isoform is predicted to terminate in exon 6. The C-terminal peptides of the truncated isoforms in the human and mouse are quite similar, supporting the idea that p19 is a functional protein variant of HRAS. We have not detected the p19 isoform by immunoblot in our system. However, p19 was detected in the nucleus and cytoplasm of HeLa cells using an antibody specifically targeting its divergent C-terminus (Guil et al., 2003a). This p19 form failed to bind to the known p21 interactors Raf1 and Rin1(Camats et al., 2009; Guil et al., 2003a). When overexpressed in different settings, p19 was found to bind a variety of proteins including p73, MDM2, Neuron-Specific Enolase, and RACK, and to have varying effects on cell growth and physiology (Camats et al., 2009; García-Cruz et al., 2015; Guil et al., 2003a; Jang et al., 2010; Jeong et al., 2006). These findings indicate that p19 likely serves a separate role from p21 and that modulation of HRAS splicing will alter p19 function in addition to changing p21 activity.

Earlier studies identified a regulatory element called rasISS1 in the intron downstream of HRAS exon 5 that acted to silence exon 5 splicing (Guil et al., 2003b). This inhibition was also observed in an in vitro splicing system, where it required the protein hnRNP A1, and was counteracted by the SR proteins, SRSF2 (SC35) and SRSF5 (SRp40). The RNA binding proteins FUS/TLS and hnRNP H, and the RNA helicase p68 (DDX5) were also found to bind rasISS1. Depletion of p68 in vivo led to increased exon 5 splicing, indicating that it might contribute to splicing repression by rasISS1. The rasISS1 was predicted to form a basepaired stem with exon 5 that may inhibit its splicing, and p68 was shown to unwind the exon 5 – rasISS1 stem in vitro. Knockdown of FUS/TLS or hnRNP H reduced the abundance of p19 HRAS protein in vivo (Camats et al., 2008). These and our findings indicate that HRAS exon 5 is modulated by a combination of positive and negative acting factors as seen with most alternative exons.

The ASO tiling approach allowed us to map regulatory elements more comprehensively across the exon 5 region. ASO’s I5-1 and I5-2 target the inhibitory sequence of rasISS1 element and the activation of splicing by I5-1 indicates the presence of a silencer. The limited effect of ASO I5-2 may come from the ASO not being able to disrupt the proposed secondary structure. Our analysis identified enhancer elements near the 3’ end of the ISS and downstream that contain G-runs, and which bind hnRNP’s H and F. We find that these proteins strongly activate splicing despite the presence of the rasISS1, and this activation requires the G-runs in the I5-3 region. The ASO tiling also indicated the presence of multiple exonic enhancers within exon 5. Some of these may recruit SRSF2 and SRSF5 whose activity on the exon was previously reported (Guil et al., 2003b). We confirmed SRSF5 as splicing activator through transient overexpression (data not shown). The ASO tiling can be refined and in future work we can more precisely delineate the cis-elements using overlapping oligos to identify those that most strongly shift exon 5 splicing and the expression of the p21 HRAS oncoprotein. These can then be tested for effects on tumor growth. Other studies examining the programs of splicing regulation in cancer have also identified HRAS as a Myc dependent exon. Expression of several SR proteins is altered in response to Myc (Anczuków et al., 2012, 2015), and it was recently reported that some SR proteins, particularly SRSF2, act to repress HRAS cassette exon splicing in Myc-active tumors (Urbanski et al., 2022).

HnRNP H has been connected to other aspects of Ras signaling. Studies of the ARaf kinase found that splicing to create its full length isoform required hnRNP H (Rauch et al., 2010, 2011). This isoform inhibits apoptosis in tumor cells through interaction with the MST2 kinase.

The short ARaf isoform expressed in low hnRNP H conditions can act as a dominant negative protein to suppress Ras activation and oncogenic transformation. These studies found that high Myc correlated with high hnRNP H expression in HeLa and several other tumor cell lines, the opposite of the correlation we observe in most tumors in the TGCA database. It will be interesting to investigate whether these different results arise for differences between cell lines and primary tumors, differences between tumor types, or some other difference between the systems. Another earlier study found that hnRNP F affected cell proliferation through interactions with mTOR and the S6 Kinase 2 pathway (Goh et al., 2010). We found that both hnRNP H and F are required for growth of a prostate cancer cell line. It will be interesting to assess the signaling pathways involved in these effects on cell proliferation control, and whether the Ras-MEK-ERK or Ras-PI3K-AKT pathways are involved.

There are several findings within our data that remain unexplained. One question is the apparent upregulation of hnRNP F by Myc. Since hnRNP F also seems to stimulate HRAS exon 5, one would expect it to go down with increased Myc. It is possible that other factors, of the many affecting exon 5, are counteracting the effect of hnRNP F. It should also be noted that Myc correlations with gene expression are only measuring RNA and it is possible the proteins encoded by these mRNAs are behaving differently. It is also possible that modifications of the hnRNP F protein could alter its activity. Another question regards the requirement for hnRNP H in the growth of Myc transformed cells. Although reduced splicing of HRAS exon 5 resulting from reduced hnRNP H is apparently conducive to growth, it some level of hnRNP H is still required. What mRNA isoforms are responsible this hnRNP H dependence will be interesting to investigate.

## MATERIALS AND METHODS

### Construction of minigene reporters and cDNA expression vectors

The HRAS minigene reporter was constructed by PCR amplifying a ∼900bp region spanning exon 4 to exon 6 from human genomic DNA isolated from HEK293 cells using Phusion high-fidelity DNA polymerase (NEB). The PCR fragment were inserted into pcDNA3.1(+) vector through restriction sites BamHI/EcoRI. G-run point mutations were introduced using site-directed mutagenesis. Construction of the C-terminal 6xHistidine tagged hnRNP H and hnRNP F cDNA constructs, and siRNA-resistant hnRNP H construct was described previously (Nazim et al., 2016). Mutations to create the siRNA-resistant hnRNP F construct were introduced by site-directed mutagenesis. hnRNP H coding sequences were subcloned into C-terminal p3xFLAG CMV vector. All the constructs were confirmed by sequencing.

### Cell culture, plasmid and siRNA transfections

HEK293, HepG2, PC3, DU145 cells were maintained in DMEM, EMEM, F12K, RPMI1640, respectively, all supplemented with 10% fetal bovine serum at 37C in 5% CO2. Cells were plated the day before transfection in a 6-well culture plate. Plasmids (3ug per well) were transfected into HEK293 cells using lipofectamine 2000 (Invitrogen), according to manufacturer’s instructions. Cells were harvested 48hrs post transfection for RT-PCR and immunoblotting analyses. siRNAs were transfected into HEK293 (10nM) or HepG2, PC3 and DU145 (20nM) using lipofectamine RNAiMAX (Invitrogen), according to manufacturer’s instructions. 24hrs after transfection, siRNAs were transfected a second time. 72hrs after the first siRNA transfection, cells were harvested for RT-PCR and immunoblotting analyses. A list of siRNA sequences is presented in the supplemental table S3.

### ASO transfections

ASOs were provided by IONIS Pharmaceuticals and have uniform phosphorothioate backbone chemistry with modified 2’ methoxyethyl sugars. HEK293 cells were plated the day before transfection in 12-well culture plates. ASOs (100nM) were transfected into cells using lipofectamine 2000 (Invitrogen). Cells were harvested 24 hrs after transfection for RT-PCR analyses. A list of ASO sequences is presented in the supplemental table S1.

### RNA isolation, RT-PCR, RT-qPCR

Total RNA was isolated using TRIzol Reagent (Life Technologies), followed by DNase I treatment (Invitrogen Ambion), and then extracted again with acid phenol chloroform (Invitrogen Ambion). 500ng – 1ug of total RNA was subjected to cDNA synthesis using SuperScript III reverse transcriptase (Thermo Fisher Scientific), primed by random hexamers. PCR was conducted using GoTaq, 18 cycles for minigene reporters and 24 cycles for endogenous RNAs. PCR products were run on 5% PAGE, stained with SYBRGold (Thermo Fisher Scientific) (unless otherwise specified), and scanned with a Typhoon imager (GE healthcare). PCR band intensities were quantified using ImageJ. RT-qPCR was conducted with SensiFast SYBR lo-ROX Kit (Bioline) on a Quant Studio 6 Flex real-time PCR system (Thermo Fisher Scientific).

RT-PCR primers are listed in the supplemental table S2.

### Immunoblotting

Proteins were extracted using RIPA lysis buffer (Thermo Fisher Scientific) supplemented with Benzonase Nuclease (Sigma Aldrich) and protease inhibitor cocktail (Sigma Aldrich). Protein concentrations were quantified by BCA protein assay kit (Pierce Biotechnology). Cell lysates were prepared with SDS loading buffer (50 mM Tris·Cl, pH 6.8, 0.05% bromophenol blue, 10% glycerol, 2% SDS, and 0.1 M DTT) and proteins were separated on SDS-PAGE gels and transferred to PVDF membranes. Antibodies used in this study were listed in the supplemental table S4. Western blot images were scanned on a Typhoon imager (GE healthcare) and band intensities were quantified using ImageJ.

### RNA Immunoprecipitation

Antibodies targeting proteins of interest or IgG isotype control (5ug) were incubated with 20ul of Dynabeads Protein G (Thermo Fisher Scientific) in buffer WB150 (20 mM HEPES-KOH pH 7.5, 150 mM NaCl, 0.1% Triton-X100) at 4C for 2 hrs. HEK293 cells were harvested and sonicated in cold lysis buffer (20 mM HEPES-KOH pH 7.5, 150 mM NaCl, 0.5% Triton X-100, 0.05% SDS, 1 mM EDTA, 0.5 mM DTT, 1x protease inhibitors, 100U/ml RNaseOUT). After centrifugation at 20,000g for 10min at 4C, the supernatant was incubated with antibody conjugated beads at 4C for 3 hrs. The beads were washed five times with buffer WB150 and RNA was extracted with TRIzol.

### Cell cycle analysis

PC3 cells were plated the day before transfection in a 6-well culture plate. siRNA were transfected as described above. 72hrs after transfection, cells were trypsinyzed and washed with PBS. These cell suspensions were fixed with 70% ice-cold ethanol overnight at -20C. Cells were stained with propidium iodide at 75 ug/ml and incubated at 37C for 30min, followed by FACS analysis.

### RNA-seq data processing, gene expression and splicing analysis for cell lines

The fastq files of HNRNPH1 and HNRNPF CRISPR RNA-seq were downloaded from ENCODE Consortium (Dunham et al., 2012) (https://www.encodeproject.org/) (HepG2 HNRNPF CRISPR KO, ENCODE accession no. ENCSR599NNK; HepG2 HNRNPH1 CRISPR KO ENCSR094HEU). The RNA-seq fastq files of Myc cell lines were downloaded from Gene Expression Omnibus (accession no. GSE141633) via fastq-dump in SRA toolkit. The raw sequencing reads were aligned to human reference genome and gene annotation (GRCh37, GENCODE release 26) using STAR (v2.7.3a) (Dobin et al., 2013). Alternative splicing in the samples was quantified as Percent Spliced In (PSI, *ψ*) using rMATS-turbo (v4.1.1) (Shen et al., 2014) by using the same gene annotation as the alignment step. Skipped exon (SE) events with insufficient average coverage, *IC* + *SC* ≤ 10, were excluded from further analysis. Significant SE events were identified using |Δ*ψ*| > 0.05 *and FDR* < 0.05, calculated using PAIRADISE (Demirdjian et al., 2020) with the equal variance option. Gene Ontology analysis was done using PANTHER (Thomas et al., 2022), p-value was computed using Fisher’s exact test, and the Benjamini-Hochberg procedure was used to control for FDR < 0.05.

### RNA-seq data compilation and processing for the pan-cancer analysis

RNA-seq data of tissue samples were compiled from two public domains: TCGA (Weinstein et al., 2013), GTEx (Lonsdale et al., 2013). Tumor tissue samples were obtained from TCGA while normal tissue was obtained from GTEx. The TCGA RNA-seq fastq files were downloaded from GDC through their GDC Data Transfer Tool Client (Grossman et al., 2016) while the GTEx files were downloaded from dbGAP (Mailman et al., 2007) through fastq-dump from the SRA toolkit. The TCGA and GTEx samples were matched by tissue origin (Robinson et al., 2019). The fastq files were mapped by STAR 2.5.3a (Dobin et al., 2013). The STAR 2-pass function was enabled to improve the accuracy of the alignment. The genome annotation file was obtained from GENCODE V26 (Harrow et al., 2012) under human genome version hg19. Gene expression quantification and alternative splicing quantification were done using Cufflinks v2.2.1 (Trapnell et al., 2012) and rMATS 4.1.0 (Shen et al., 2014) respectively. Percent Spliced In (PSI) ratio was used as a statistic to quantify alternative splicing events from the rMATS output. Splicing events were filtered for splice junction reads >= 10 (otherwise the sample’s PSI value is marked with ‘NA’), PSI range (i.e. difference between maximum and minimum PSI values) > 5%, and for mean skipping or inclusion values over 5% across the entire dataset, with fewer than 50% of samples missing values.

### Pathway Enrichment-Guided Activity Study of Alternative Splicing for the pan-cancer analysis

The PEGASAS pipeline (Phillips et al., 2020) was used to calculate pathway activity scores and identify alternative splicing events that correlate with the signaling pathway of interest. The Myc Targets V2 hallmark gene signature list obtained from The Molecular Signatures Database (MSigDB) (Liberzon et al., 2011) was used for pathway activity score calculation. Myc pathway scores were calculated using cufflinks expression outputs. Alternative splicing events (SE) correlated with the Myc signaling pathway were identified through the PEGASAS correlation step. A correlation permutation test was used to acquire empirical p-values, by permuting the pathway scores 5,000 times. Highly correlated events were defined as events with an empirical p-value < 0.0002 and with a |Pearson correlation coefficient| > 0.3, as in previous study (Phillips et al., 2020).

### Gene expression values for RBP correlation

FeatureCounts v2.0.1 was used to quantify reads from aligned BAM files (Liao et al., 2014). DESeq2 v1.26.0 was used to normalize gene expression (Love et al., 2014). The normalized gene expression values for HNRNPH1 and HNRNPF along with 218 other splicing factors were correlated with Myc scores (Han et al., 2013).

## ACKNOWLEDGEMENTS

The other authors dedicate this report to the late John W. Phillips. We thank Nazim Mohammad for the hnRNP H/F plasmids; Liang Wang (the Witte Lab) for the prostate cancer cell lines; all members of the Black laboratory for help and discussions. This work was supported by funds from National Institutes of Health (NIH) R01CA220238 to O.N.W., Y.X., and D.L.B.; U01 CA- 233074-01 to O.N.W. and Y.X.; R35GM136426 to D.L.B; R01GM088342 and R56HG012310 to Y.X.; A Directors Award from the Jonsson Comprehensive Cancer Center at UCLA to D.L.B.; and Developmental Research Program Awards from the UCLA SPORE in Prostate Cancer (Award P50 CA092131) to J.W.P. and D.L.B.. X.C. was supported by Whitcome and Warsaw family fellowships at UCLA. J.W.P. was supported by a postdoctoral fellowship from training grant DOD W81XWH-16-1-0216 at UCLA. The results shown here are in part based upon data generated by the TCGA Research Network (https://www.cancer.gov/tcga), the Genotype-Tissue Expression (GTEx) Project (https://www.commonfund.nih.gov/gtex), and the ENCODE Consortium (https://www.encodeproject.org/).

## Author Contributions

X.C., J.W.P, O.N.W., Y.X., and D.L.B. designed the research; X.C. performed experiments and analyzed data; X.C., H.T.Y., and B.Z. performed bioinformatic analysis; J.W.P, D.C. and F.R. contributed analytic methods/reagents; and X.C. and D.L.B wrote the manuscript with input from the other authors.

---

**Figure S1.**
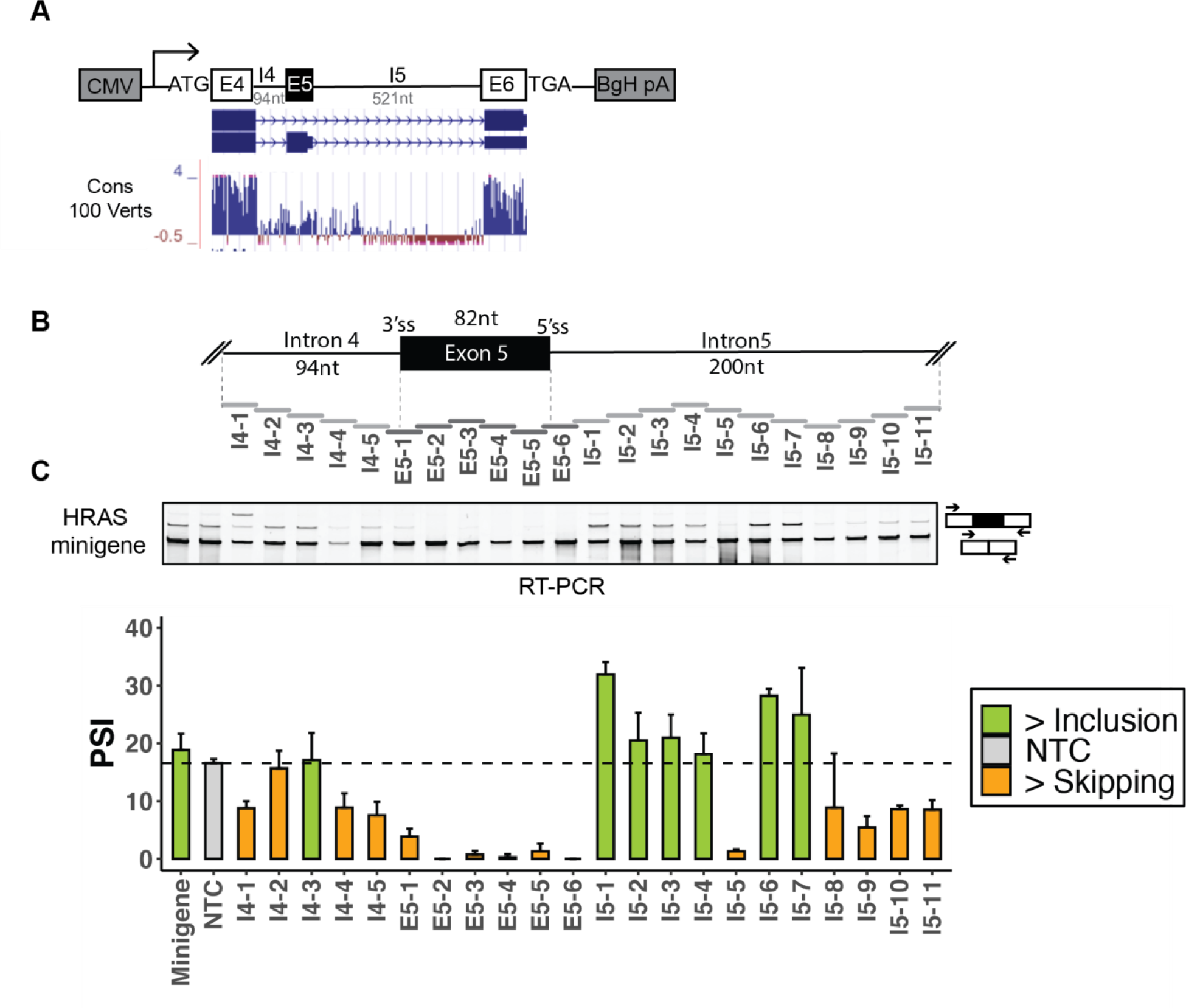
Identification of splicing cis-regulatory elements with HRAS minigene reporter. **(A)** Diagram of HRAS minigene reporter with a schematic representation of gene structure and conservation. **(B)** Schematic of ASO tiling across HRAS cassette exon and its partial flanking introns on minigene reporter. A 376-nucleotide region tiled by non-overlapping 18-mer ASOs. Each horizontal bar represents an ASO. (C) Semi-quantitative RT-PCR analyses showing the effects of ASOs on HRAS minigene reporter. The arrows indicate the locations of minigene-specific primers. The bar graph shows the quantification of RT-PCR results calculated as percent-spliced-in (PSI) (gray: control; orange: more skipping; green: more inclusion). Each bar shows the mean PSI +/- SD of three biological replicates. NTC, non-targeting control. Minigene, no-ASO mock control.

**Figure S2.**
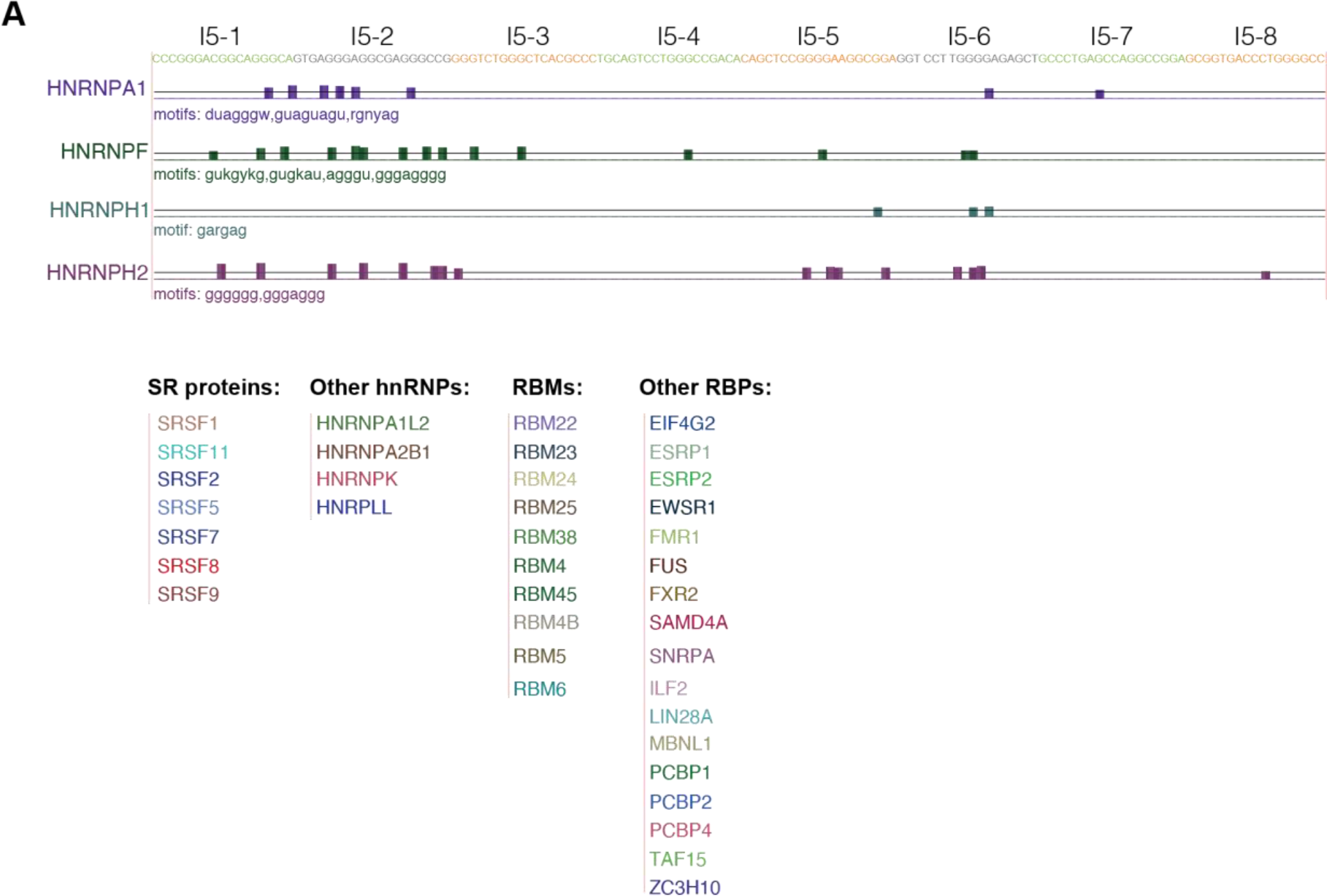
RBPmap identifies binding motifs of potential splicing regulators in the HRAS intron 5 sequence. **(A)** UCSC genome browser visualization of representative RBPmap- predicted splicing factor binding sites on HRAS intron 5. The sequences targeted by each ASO are highlighted in orange (more skipping), green (more inclusion), gray (no change). Below are lists summarizing other potential splicing factors identified by the program, including SR proteins, other hnRNPs, RBMs, and others. Note that the motif listed for hnRNP H in this analyses is only one of a set of known hnRNP H motifs, which also include those listed for hnRNP F (Dominguez et al., 2018; Huelga et al., 2012; Uren et al., 2016).

**Figure S3.**
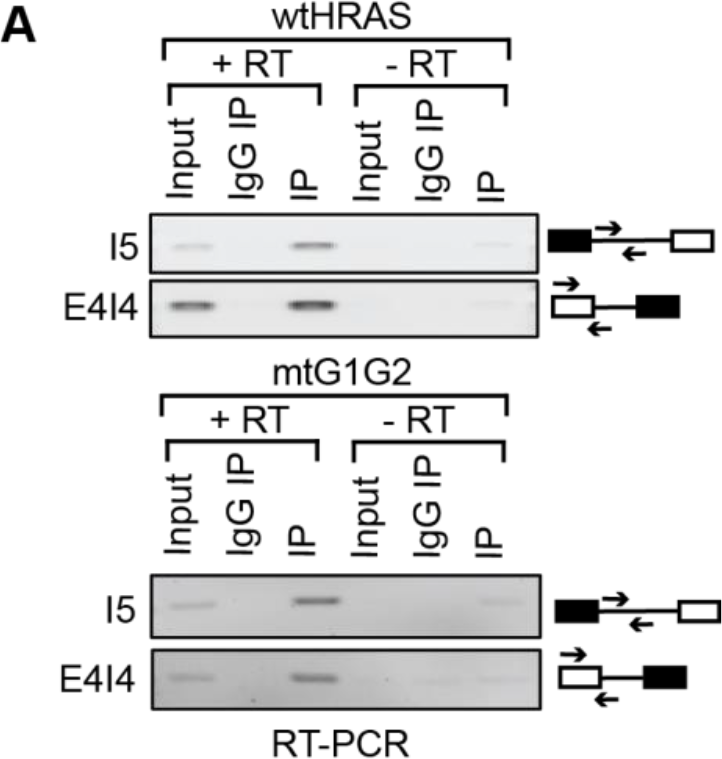
Immunoprecipitated RNA is free of minigene DNA contamination. **(A)** Standard RT-PCR of immunoprecipitated RNA using primer pairs amplifying a segment of intron5 or the exon4-intron4 junction, in the presence [+] or absence [-] of reverse transcriptase. Immunoprecipitated RNA (80%) was amplified with 5 more PCR cycles than the Input control (5%). wt, wild type; mt, mutant.

**Figure S4.**
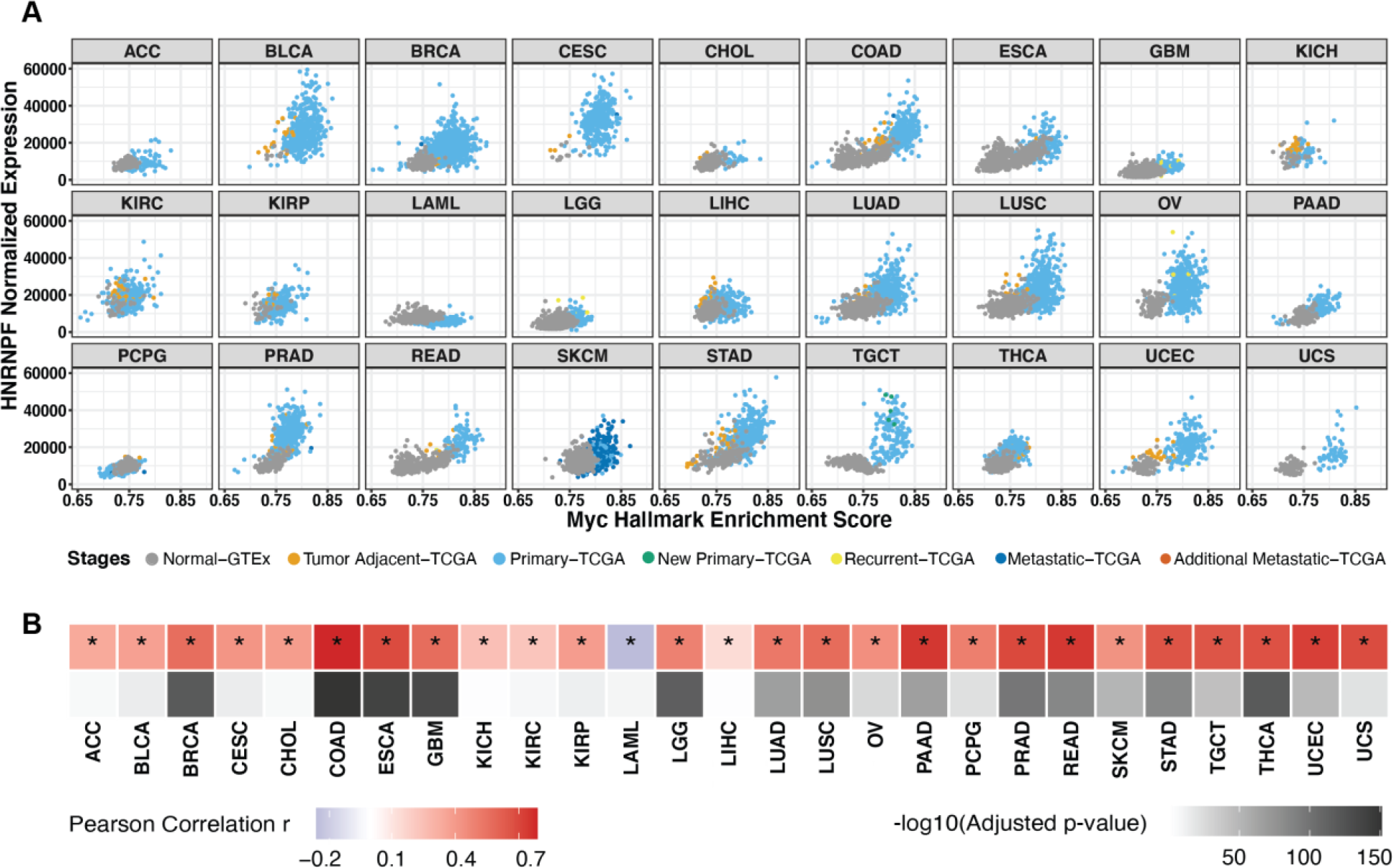
HNRNPF gene expression is upregulated with Myc activation across tumor types. **(A)** Scatterplot matrix showing the correlation of normalized HNRNPF RNA level versus Myc hallmark enrichment score spanning the disease spectrum in multiple tumor types. **(B)** Heatmap summarizing the Pearson correlation coefficient for each tumor type, accompanied with its corresponding adjusted p-value. *, tumor types with statistically significant adjusted p-value (< 0.05).

**Figure S5.**
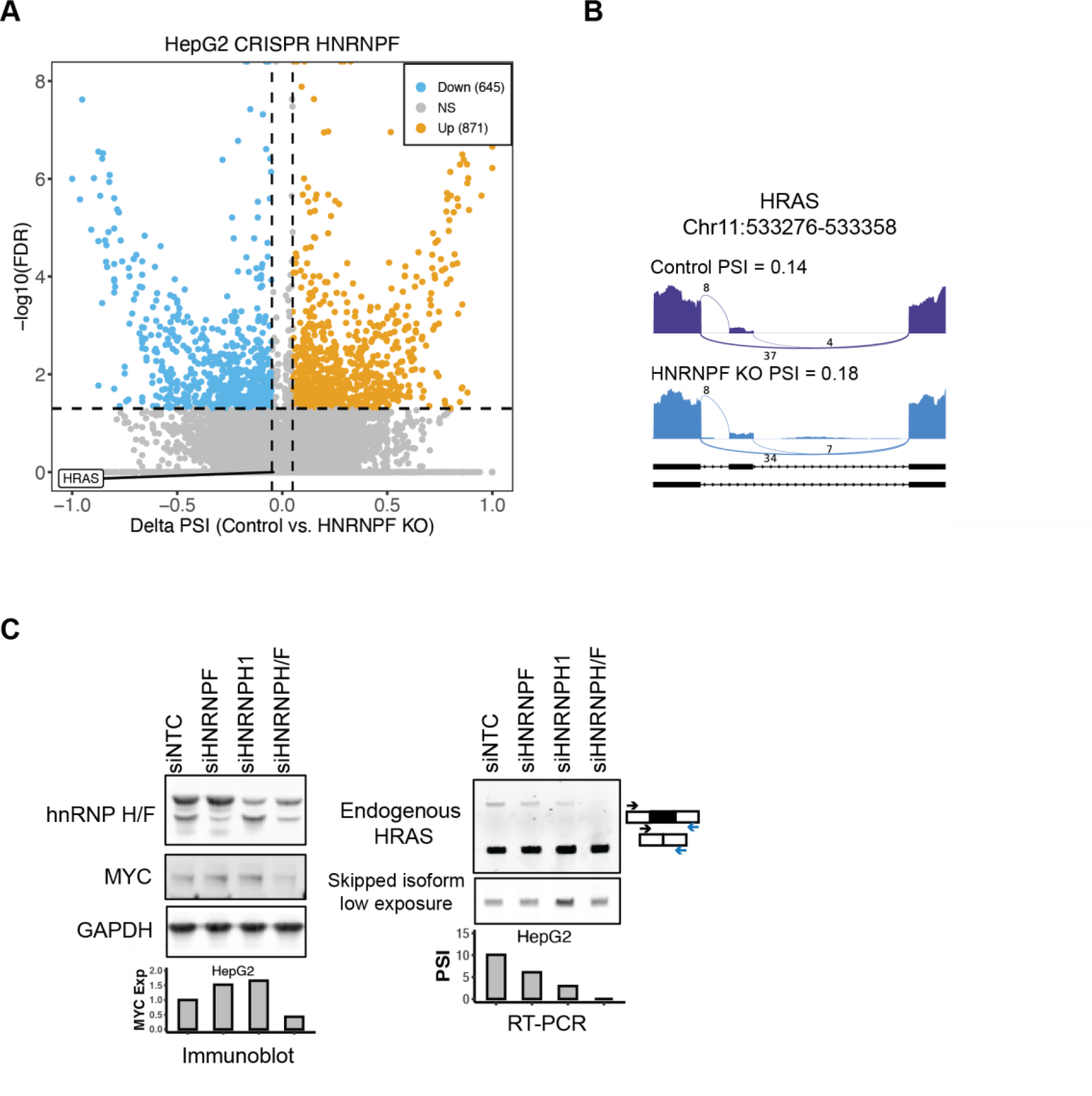
Differential splicing analysis after HNRNPF depletion from HepG2 cells. **(A)** Scatterplot showing the SE events detected after HNRNPF CRISPR knockout in HepG2 cells compared to control. RNA-seq data from ENCODE was analyzed. Significant SE events were filtered by junction reads per event > 10, |deltaPSI| > 0.05, and FDR < 0.05. **(B)** Sashimi plot showing the PSI of HRAS exon 5 in hnRNPF KO and control cells. KO, knockout; NS, non-significant. **(C)** The effects of hnRNP H/F knockdown on HRAS splicing in HepG2 cells. Immunoblot showing the protein levels of hnRNP H/F and MYC in HepG2 cells after transfection with control, HNRNPF, HNRNPH1, or HNRNPH/F siRNAs. GAPDH was used as a loading control. The bar graph (left) shows the quantification of MYC expression in response to each siRNA perturbation. RT-PCR analysis of the splicing of endogenous HRAS transcripts after siRNA treatment. The bar graph (right) shows the quantification of HRAS PSI.

**Table S1.**
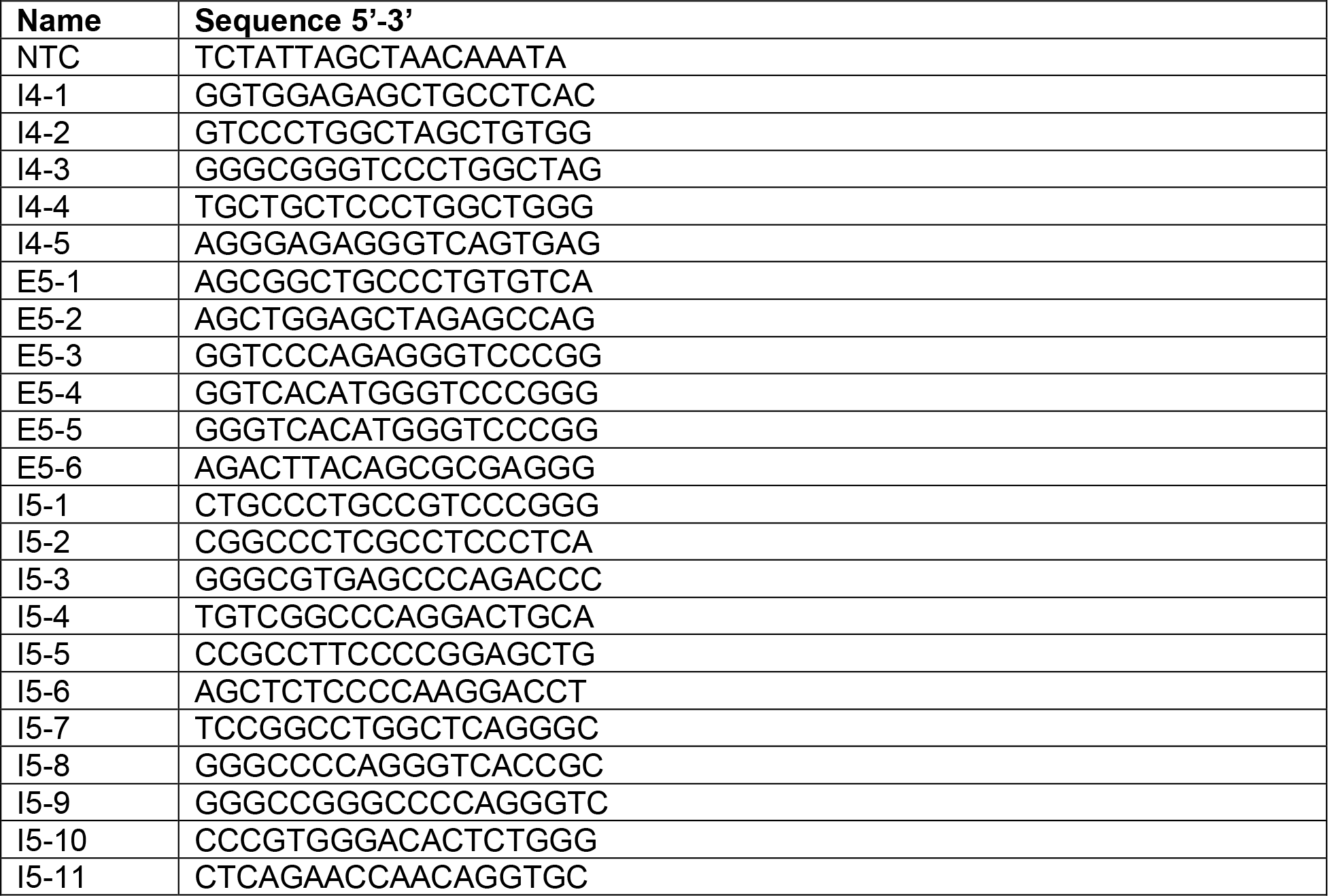
List of ASOs

**Table S2.**
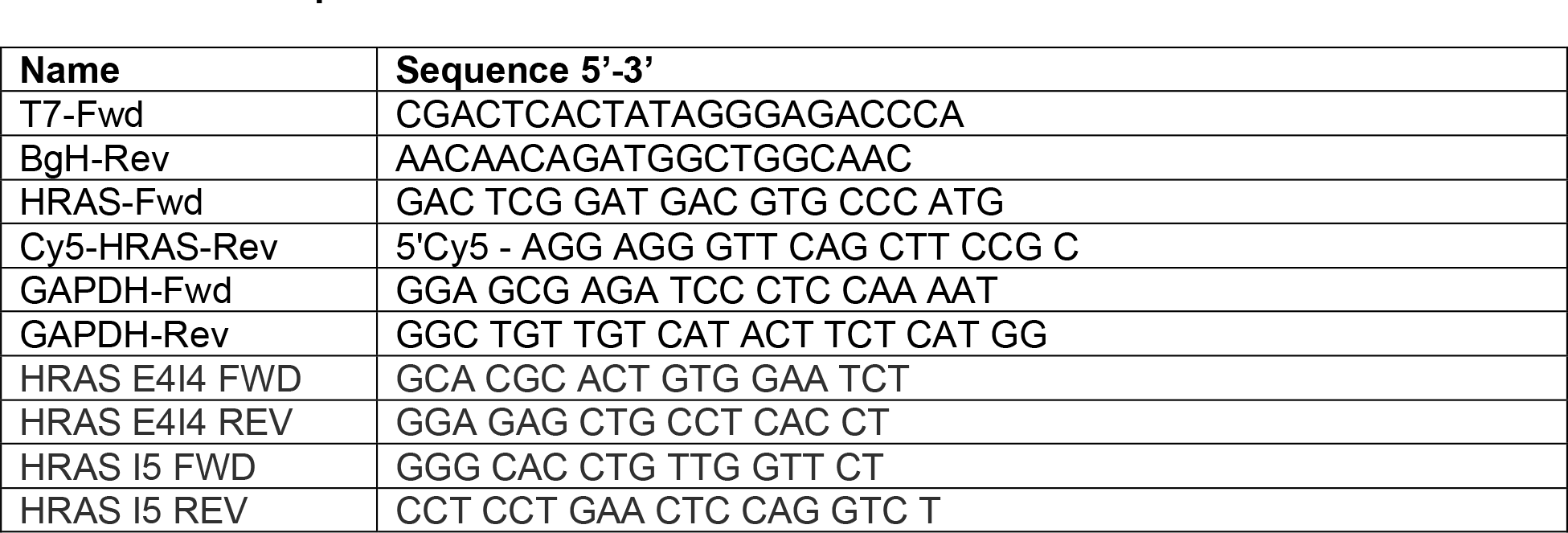
Lists of primers

**Table S3.**
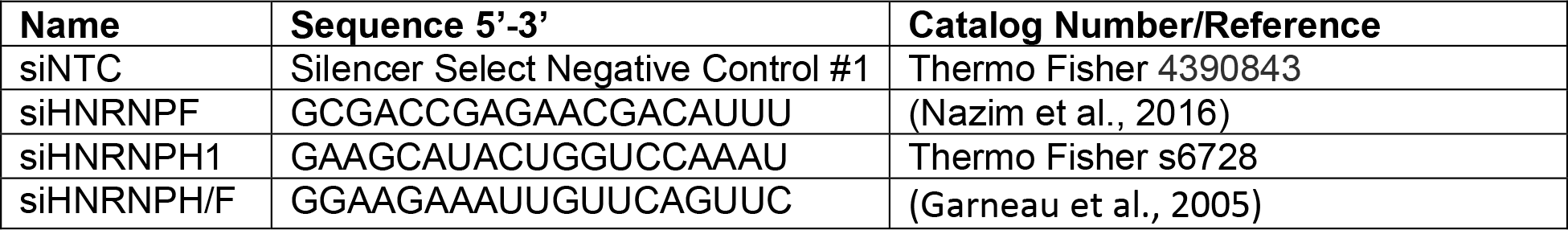
**Lists of siRNAs**

**Table S4.** Lists of antibodies

| **Name** | **Manufacturer** | **Catalog Number** |
| --- | --- | --- |
| Anti-hnRNP H | Lab made | N/A |
| Anti-hnRNP F | Lab made | N/A |
| Anti-hnRNP H/F | Santa Cruz Biotechnology | sc-32310 |
| Anti-His | MBL International | D291-3 |
| Anti-GAPDH | Santa Cruz Biotechnology | sc-47724 |
| Anti-HRAS | Abcam | ab86696 |
| Anti-Flag | Sigma Aldrich | F3165 |
| Mouse IgG isotype control | Invitrogen | 14471485 |
| Rabbit IgG isotype control | Thermo Fisher | 02-6102 |
| Anti-hnRNP H (IP) | Abcam | ab10374 |
| Anti-hnRNP F (IP) | Sigma Aldrich | 04-1462 |
| Anti-MYC | Abcam | ab32072 |
| Anti-cPARP | Cell signaling | 5625S |

**Table S5.** Splicing changes of other G-run mutations

| **ASO Target** | **Mutations** | **Splicing changes** | **Consistency with ASO hits** |
| --- | --- | --- | --- |
| I5-1 first G-run | ggg to gcg | No change | NA |
| I5-1 second G-run | ggg to gcg | Exon skipping | No |
| I5-4 | ggg to gcg | Slight exon inclusion | Yes |
| I5-6 | gggg to gctg | Exon inclusion | Yes with minigene data; No with endogenous data |

